# Revisiting the source of wilt symptoms: X-ray microcomputed tomography provides direct evidence that *Ralstonia* biomass clogs xylem vessels

**DOI:** 10.1101/2021.03.19.436187

**Authors:** Brian Ingel, Denise Caldwell, Fiona Duong, Dilworth Y. Parkinson, Katherine A. McCulloh, Anjali S. Iyer-Pascuzzi, Andrew J. McElrone, Tiffany M. Lowe-Power

## Abstract

Plant pathogenic *Ralstonia* cause wilt diseases by colonizing xylem vessels and disrupting water transport. Due to the abundance of *Ralstonia* cells in vessels, the dogma is that bacterial biomass clogs vessels and reduces the flow of xylem sap. However, the physiological mechanism of xylem disruption during bacterial wilt disease is untested. Using a tomato and *Ralstonia pseudosolanacearum* GMI1000 model, we visualized and quantified the spatiotemporal dynamics of xylem disruption during bacterial wilt disease. First, we measured stomatal conductance of leaflets on mock-inoculated and wilt-symptomatic plants. Wilted leaflets had reduced stomatal conductance, as did turgid leaflets located on the same petiole as wilted leaflets. Next, we used X-ray microcomputed tomography (X-ray microCT) and light microscopy to differentiate between mechanisms of xylem disruption: blockage by bacterial biomass, blockage by vascular tyloses, or sap displacement by gas embolisms. We imaged stems on plants with intact roots and leaves to quantify embolized vessels. Embolized vessels were rare, but there was a slight trend of increased vessel embolisms in infected plants with low bacterial population sizes. To test the hypothesis that vessels are clogged during bacterial wilt, we imaged excised stems after allowing the sap to evaporate during a brief dehydration. Most xylem vessels in mock-infected plants emptied their contents after excision, but non-conductive clogged vessels were abundant in infected plants by 2 days post infection. At wilt onset when bacterial populations exceeded 5×10^8^ cfu/g stem tissue, approximately half of the xylem vessels were clogged with electron-dense bacterial biomass. We found no evidence of tyloses in the X-ray microCT reconstructions or light microscopy on the preserved stems. Bacterial blockage of vessels appears to be the principal cause of vascular disruption during *Ralstonia* wilt.

## Introduction

Wilt diseases are destructive plant diseases that compromise xylem function and typically kill susceptible hosts (Yadeta and Thomma 2013; Lowe-Power et al. 2018). Wilt pathogens can disrupt xylem function by several potential mechanisms (Lowe-Power et al. 2018). First, pathogens can directly clog vessels with their biomass. Second, plant defenses can block vessels. The living parenchyma cells adjacent to xylem vessels can grow balloon-shaped structures called “tyloses” that obstruct the vessels. Tyloses can completely seal off xylem vessels in response to pathogen invasion or gas embolisms (De Micco et al. 2016). Additionally vascular parenchyma cells can secrete pectic polymers that form gels that impede pathogen and sap movement (De Micco et al. 2016). While tyloses and gels can prevent the systemic spread of some pathogens, they are not effective at containing all pathogens and can cause collateral damage to the host by reducing the water transport capacity of the xylem (Yadeta and Thomma 2013; Ingel et al. 2020; Kashyap et al. 2020). Finally, vascular pathogens often secrete cellulases and pectinases (Schwarze and Landmesser 2000; Liu et al. 2005; Sun et al. 2011; Gluck-Thaler et al. 2020). These enzymes enable pathogens to move from vessel to vessel because the enzymes degrade the pit membranes that connect vessels to each other (Pérez-Donoso et al. 2010). Pit membranes are semi-permeable barriers composed of cellulose, hemicellulose, and pectin, which are more digestible polymers than the polyphenolic lignin that reinforces vessel walls (Kaack et al. 2019). In addition to blocking large particles from moving vessel-to-vessel, pit membranes also protect against the spread of gas embolisms (Tyree and Sperry 1989; Tyree and Zimmermann 2002; Choat et al. 2008). The narrow pores in pit membranes may allow the cohesive forces of water molecules to resist the entry of gases into sap-filled vessels (Sperry and Hacke 2004). Therefore, pathogen degradation of pit membranes may increase the xylem’s vulnerability to embolism propagation. Increased incidence of vessel embolism has been reported for oak trees infected with the xylem-limited bacterial pathogen *Xylella fastidiosa* (McElrone et al. 2008).

Bacterial pathogens in the *Ralstonia* species complex (*R. solanacearum, R. pseudosolanacearum*, and *R. syzygii*) cause wilt diseases on a broad range of plants in the tropics and subtropics (Lowe-Power and Chipman 2020). *Ralstonia* are soil-borne bacteria that invade plant roots and colonize the xylem before fatally wilting susceptible hosts (Caldwell et al. 2017). *Ralstonia* infection compromises the xylem’s ability to transport water; before onset of wilt symptoms, plants have a decreased rate of water uptake from the soil (Denny et al. 1990). Light microscopy and histology show that tyloses are rare in *Ralstonia*-infected plants (Vasse et al. 1995; Rahman et al. 1999; Caldwell et al. 2017). Thus, the major cause of xylem disruption does not appear to be tyloses. In contrast, microscopy of infected roots and stems shows that *Ralstonia* grows extensively in the xylem and that bacterial cells and biofilm matrix are abundant within individual vessels (Caldwell et al. 2017). Furthermore, high populations of *Ralstonia* in stems (exceeding 10^9^ cfu/g tissue) are correlated with onset of wilt symptoms (Lowe-Power et al. 2018). Thus, existing data is consistent with a model that *Ralstonia* wilt reduces sap flow by clogging xylem vessels. However, this model has not been directly tested. As a counter-point, the biofilm matrix of *Ralstonia* is highly fluidal (Dalsing and Allen 2014). A diagnostic sign of bacterial wilt disease is “bacterial streaming”, where a bacterial ooze streams out from cut stems when suspended in water (García et al. 2019). Further, *Ralstonia* biomass exudes with xylem sap when partially wilted tomato plants are de-topped. Therefore, it is possible that *Ralstonia* biomass is present in xylem vessels but does not impede flow of sap.

Similar to other xylem-inhabiting bacteria (e.g. *Xylella fastidiosa*; (McElrone et al. 2008)), we wondered whether *Ralstonia* may increase the incidence of xylem vessel embolism in the early stages of disease progression. Like *Xylella, Ralstonia* degrade xylem pit membranes (Grimault et al. 1994; Rahman et al. 1999; Liu et al. 2005; Roper et al. 2007; Ingel et al. 2019). Additionally, most *Ralstonia* are facultative anaerobes that respire nitrate in the low-oxygen xylem (Dalsing et al. 2015; Prior et al. 2016). Nitrate respiration yields N_2_ gas, which has low aqueous solubility and may form gas bubbles that embolize vessels (Weiss 1970). Destructive methods like most microscopy assays, are not able to distinguish between air-filled vessels and sap-filled vessels.

Here, we use X-ray microCT to directly investigate the mechanism of xylem disruption during *Ralstonia* wilt of tomato. X-ray microCT yields 3-D views of tissue density at the micron resolution (McElrone et al. 2013). Because X-ray microCT imaging does not require destructive sample preparation, it is well-suited to visualizing air-filled compartments in biological tissues. We coupled functional assays with X-ray microCT imaging to visualize the functionality of individual xylem vessels. We used microCT microscopy, and quantitative PCR (ddPCR) to visualize and quantify bacterial populations in xylem vessels of *Ralstonia*-infected plants. Overall, our data support the model that *Ralstonia* clogs xylem vessels during wilt disease.

## Results

### Tomato xylem function progressively decreases as wilt symptoms become more severe

Bacterial wilt disease of tomato is characterized by a progressive and rapid wilting of leaflets over a period of days to weeks. At wilt onset, plants have spatial heterogeneity in the turgidity of leaflets. We hypothesized that there is a correlation between vascular decline and symptom onset. To characterize the water status of leaflets, we quantified stomatal conductance in individual leaflets of mock-inoculated and wilt-symptomatic plants. Stomatal aperture, which largely determines stomatal conductance, is actively regulated based on water status of the leaf and is a reliable indirect metric of the functional status of the xylem vessels that supply each leaf with water (Brodribb et al. 2017).

Susceptible tomato plants were inoculated via soil-drench with *Ralstonia pseudosolanacearum* GMI1000 or mock inoculated with water. At symptom onset (3-4 dpi), stomatal conductance was measured (Fig 1). On wilt-symptomatic plants, leaflets were classified as “wilted”, “turgid near wilted” (i.e. turgid leaflets located on petioles with wilted leaflets), or “turgid” if located on petioles without symptoms (Fig 1A). Stomatal conductance of leaflets was correlated with wilt severity (Fig 1B). Turgid leaflets on mock-inoculated plants had the highest stomatal conductance (median: 294 mmol m^-2^ s^-1^)., and wilted leaflets had the lowest stomatal conductance (median: 12 mmol m^-2^ s^-1^). For turgid leaflets on symptomatic plants, there was a trend of lower stomatal conductance in leaflets that were co-located on petioles with wilted leaflets. Overall, stomatal conductance of leaflets matched the spatial heterogeneity in wilt symptoms.

**Fig 1.**
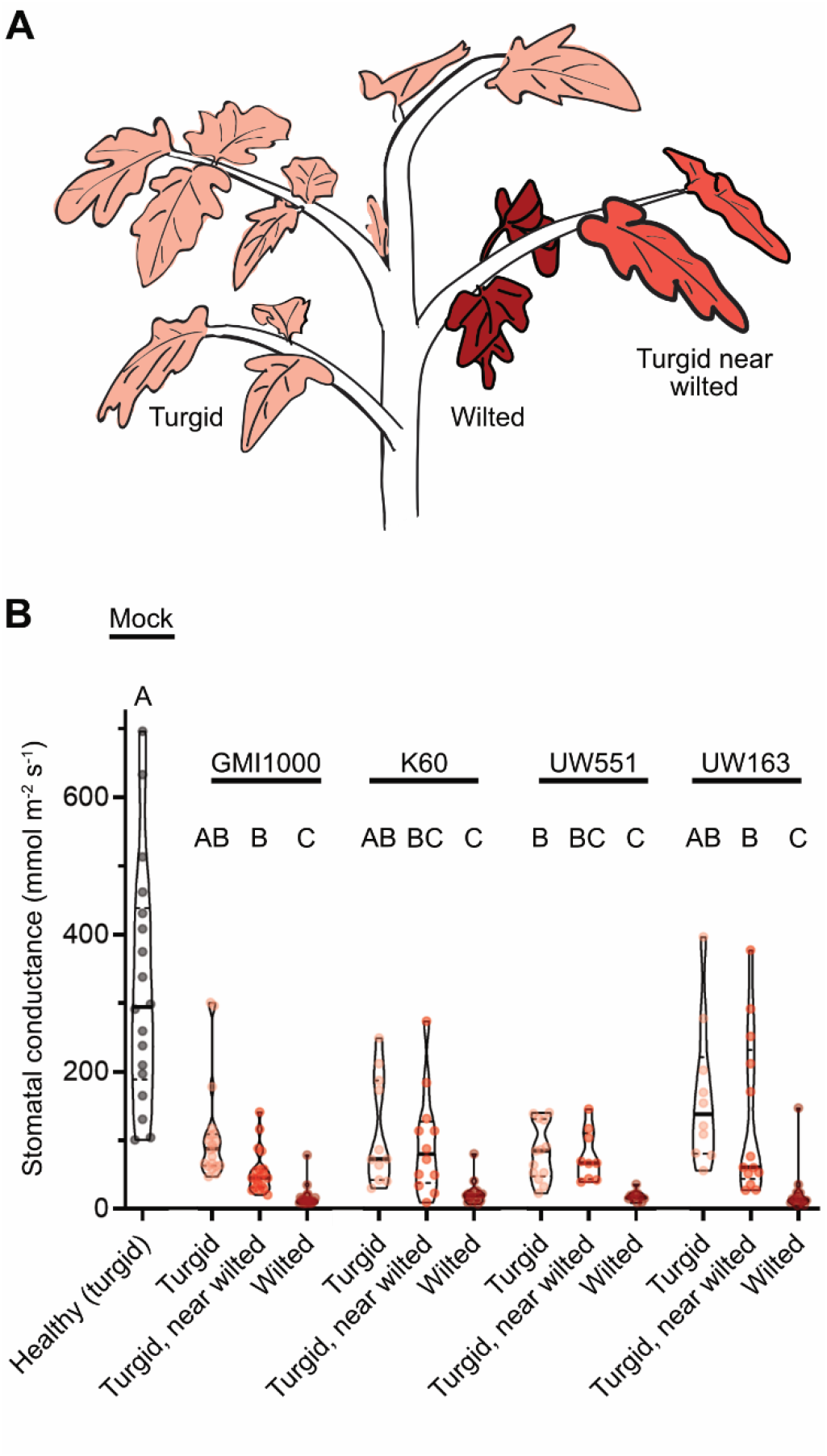
Spatial heterogeneity in leaflet wilt status reflects a stepwise decline in stomatal conductance. Plants were soil-drench inoculated with either *R. pseudosolanaceaurm* GMI1000, *R. solanacearum* K60, *R. solanacearum* UW551, *R. solanacearum* UW163, or mock-inoculated with water. Stomatal conductance of leaflets was measured with a Licor6400XT gas analyzer. Infected plants were sampled on the first day of wilt onset (3-4 days after infection). (A) Leaflets on the infected plant were categorized as turgid, wilted, or “turgid near wilted” if visually turgid but on the same petiole as wilted leaflets. (B) Stomatal conductance of leaflets on mock-inoculated or partially wilted *Ralstonia*-infected tomato plants. At least 3 leaflets of 6-10 plants were sampled per condition. Statistics: Letters indicate significance of pairwise comparisons using ANOVA with Tukey’s multiple comparison’s test.

We hypothesized that progressive declines in leaflet stomatal conductance was a general characteristic of bacterial wilt disease of tomato. We tested this by infecting tomato plants with three additional *Ralstonia* strains from different sequevar clades: *R. solanacearum* K60 (IIA-7), *R. solanacearum* UW163 (IIB-4), and *R. solanacearum* UW551 (IIB-1). Tomato plants infected with the *R. solanacearum* strains showed similar results as plants infected with *R. pseudosolanacearum* GMI1000 (I-18) (Fig 1B). These results suggested that the step-wise disruption of functional tomato xylem vessels is a common occurrence in *Ralstonia*-mediated bacterial wilt of tomato and not limited to the I-18 clade.

### Gas embolisms are rare in *Ralstonia*-infected tomato xylem

X-ray microCT allows non-destructive 3-D imaging of the density of biological tissues. We used X-ray microCT to visualize xylem vessels in stems of intact plants. We imaged the stems of mock-inoculated tomato plants and plants infected with *R. pseudosolanacearum* GMI1000 while the plants rotated 180° in an X-ray beam. We reconstructed the samples in 3-D and investigated the lumina of xylem vessels. The 11 plants each had over 50 total xylem vessels (Fig S11A). The xylem vessels were outlined by dense (whiter) rings of lignified tissue, and the lumina of most vessels were grey, indicating they were filled with xylem sap. A few vessel lumina were filled with gas embolisms (Fig 2 and Fig S2), which were black due to the lower density of gas than the density of hydrated plant tissue. One of three mock-inoculated plants had two embolized vessels, while the other two had none. Four of seven *Ralstonia*-infected plants had two-to-six embolized vessels (Fig 2C and S2). The four infected plants with embolized vessels had slightly lower bacterial burdens (1.1×10^8^ −3.8×10^8^ cfu/g) than the three infected plants without embolized vessels (4.6×10^8^ −4.7×10^9^ cfu/g).

**Fig 2.**
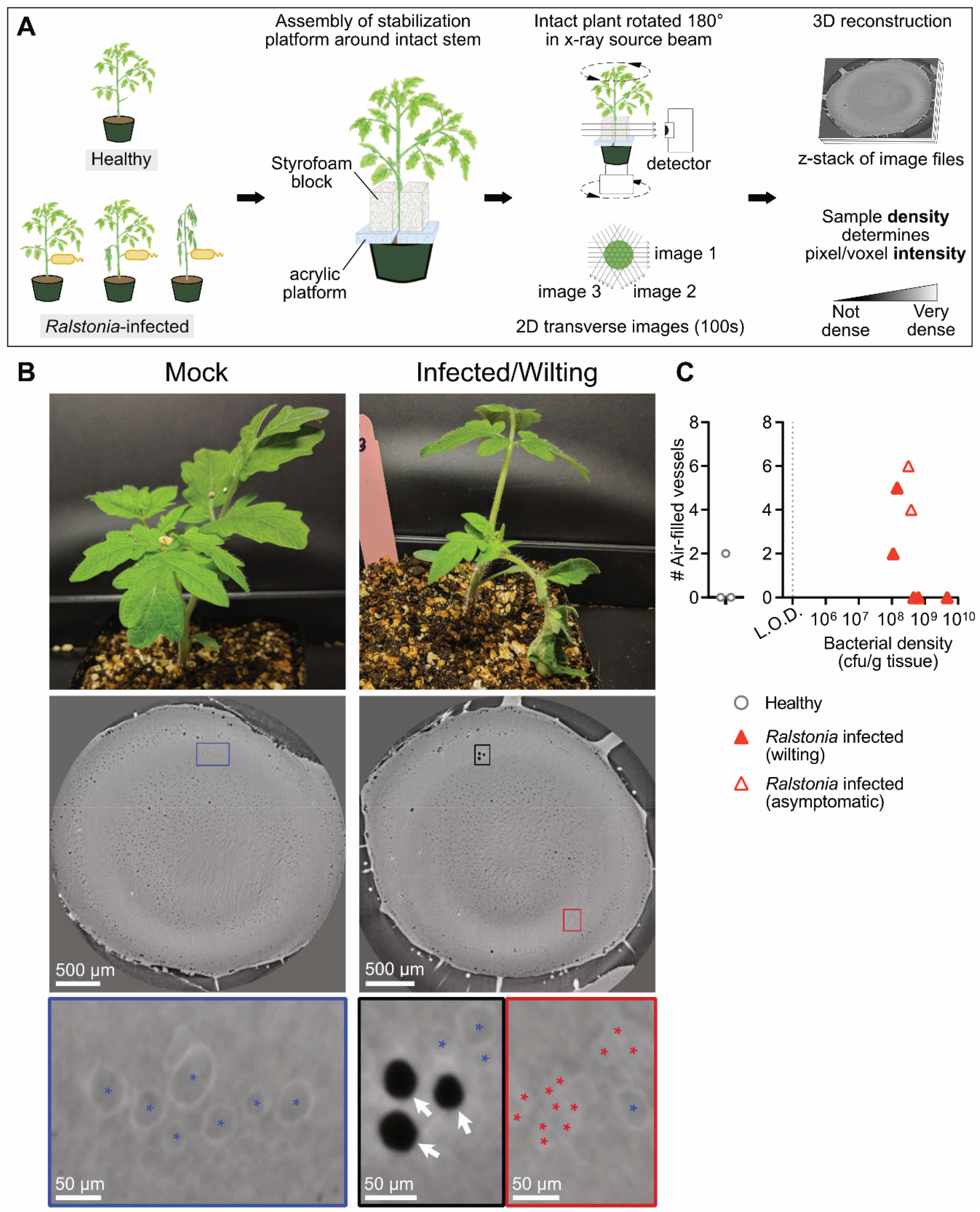
X-ray MicroCT imaging of tomato stem on intact plants reveals bacterial wilt disease increases density of vessel lumina. The stem xylem of tomato plants was inoculated with water or 10^4^ cfu *R. pseudosolanacearum* GMI1000 via a cut petiole inoculation. Stems on plants with intact root and leaf systems were imaged via X-ray microCT (N= 3 mock-inoculated plants and N=7 *Ralstonia*-infected pants). (A) Overview of experiment. (B) Representative images of whole plants and X-ray reconstructions of stem cross-sections of a mock-inoculated and *Ralstonia*-infected plant. Cross sections are located near where the inoculated petiole connects to the main stem. Vascular bundles in the boxed regions are shown at higher resolution in bottom row. White arrows point to air-filled (embolized) vessels. Colored asterisks (*) indicate vessels filled with material of varying density. Blue asterisks: grey lumina. Red asterisks: lumen content with a high density similar to the lignified vessel wall. Fig S1 and S2 include whole-plant photographs and microCT reconstructions from all samples. Fig S12 plots the relative pixel intensity of clogged, sap-filled, and empty xylem vessels. (C) The number of air-filled (embolized) vessels in stems with varying bacterial population density. Bacterial density was quantified with digital droplet PCR. The dashed grey line shows the limit of detection (L.O.D.).

Close inspection of the vascular bundles revealed that wilt-symptomatic plants had many xylem vessels with lumina that were filled with a substance denser than xylem sap and similar in density to the lignified wall (Fig 2B, red asterisks; Fig S12). In contrast, vessels in mock-inoculated plants were had a white, dense ring (lignin) that surrounded grey lumina (Fig S12). We hypothesized that the dense occlusions in wilt-symptomatic vessels were bacterial biomass clogs within vessels.

### Dense physical obstructions clog tomato xylem vessels

To test the hypothesis that bacterial biomass clogs xylem vessels and inhibits sap transport, we coupled a functional assay to the microCT imaging. To determine how many vessels were capable of sap transport, we excised stems from plants and briefly dehydrated them before X-ray imaging (Fig 3). During the dehydration, xylem sap would evaporate from the unclogged xylem vessels, but the bacterial polysaccharides from clogged vessels would resist desiccation, and remain in the xylem vessel. To synchronize infection, plants were inoculated by a cut-petiole method where 10^4^ cfu or 10^6^ cfu *R. pseudosolanacearum* GMI1000 (or water) was directly introduced into the stem xylem. In the first trial, we imaged stems from mock-inoculated plants and infected plants 1 day post inoculation (dpi) and 2 dpi with 10^4^ cfu. In the second trial, we imaged stems from the same conditions and added an additional condition where plants were inoculated with a higher dose of *Ralstonia* (10^6^ cfu) and imaged at 2 dpi.

**Fig 3.**
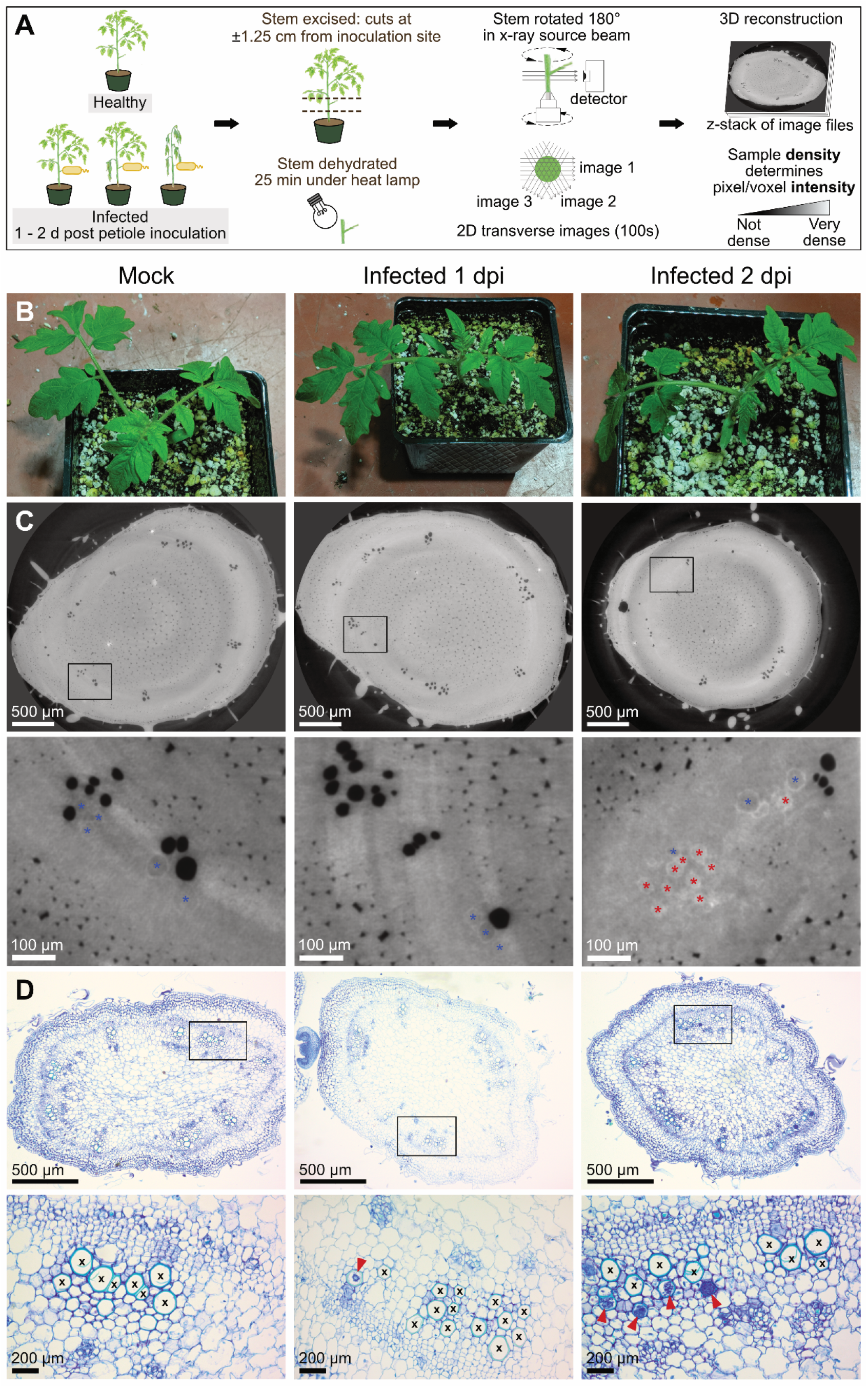
MicroCT imaging of excised, dehydrated tomato stems reveals *Ralstonia* clogs vessels and reduces xylem vessel function. Tomato plants were inoculated with water or *R. pseudosolanacearum* GMI1000 (10^4^ cfu or 10^6^ cfu) via a cut petiole inoculation and were imaged at 1-2 dpi. Before imaging with X-ray microCT, 2 cm segments of stem around the inoculation site were excised and dehydrated briefly to allow sap to evaporate from functioning xylem vessels. Two trials were performed with N ≥ 5 plants per condition. (A) Experimental overview. (B-D) Representative photographs of whole plants, microCT reconstructions of excised stem, and histology micrographs of preserved stem. Full images are available in Fig S3-S8. (C) Images show microCT cross-sections of stem near the petiole inoculation site. Black boxes are shown with enhanced resolution below each cross-section. Symbols indicate vessels filled with material of varying density. Blue asterisks: grey lumina with low-density contents. Red asterisks: lumen content with a high density similar to the lignified vessel wall. Fig S12 plots the relative pixel intensity of clogged, sap-filled, and empty xylem vessels. (D) Preserved stems were stained with toluidine blue and imaged using light microscopy (4X magnification; top row). Black boxes are shown with enhanced magnification (20X; bottom row). Black “X” are displayed in open vessels free of occlusions. Red arrows indicate vessels partially or fully occluded by *R. pseudosolanacearum* GMI1000.

We quantified the number of sap-conducting vessels from microCT cross-sections (Figs 3 and S3-S8; Fig S11B; Fig 4A). As expected, the majority of xylem vessels in mock-inoculated plants released their contents during dehydration. The reconstructions showed numerous black, empty vessels that were clustered in vascular bundles around the periphery of the stem (Fig 3C). In trial 1, mock-inoculated plants had 59-81 functional vessels (Fig 4A and S5). In the second trial, healthy stems had 114-133 functional vessels because the plants grew faster prior to inoculation (Fig 4A and S5). Stems from the 1 dpi infected plants and the mock-inoculated plants had similar numbers and distributions of sap-conducting vessels in both trials (Fig 4A and S6). By 2 dpi, infected plants (both asymptomatic and symptomatic) had fewer black, empty vessels in both trials, indicating fewer sap conducting vessels (P<0.05 Kruskal-Wallis test with Dunn’s multiple comparison’s test) (Fig 4A and S7). In trial 2, the 2 dpi plants inoculated with the high titer of bacteria (10^6^ cfu) had fewer sap conducting vessels than the 2 dpi plants inoculated with the moderate 10^4^ cfu titer (Fig 4A, S7, and S8). Furthermore, many of the 2 dpi stems from infected plants in both trials had large regions that lacked any sap-conducting vessels (Fig S7-S8).

**Fig 4.**
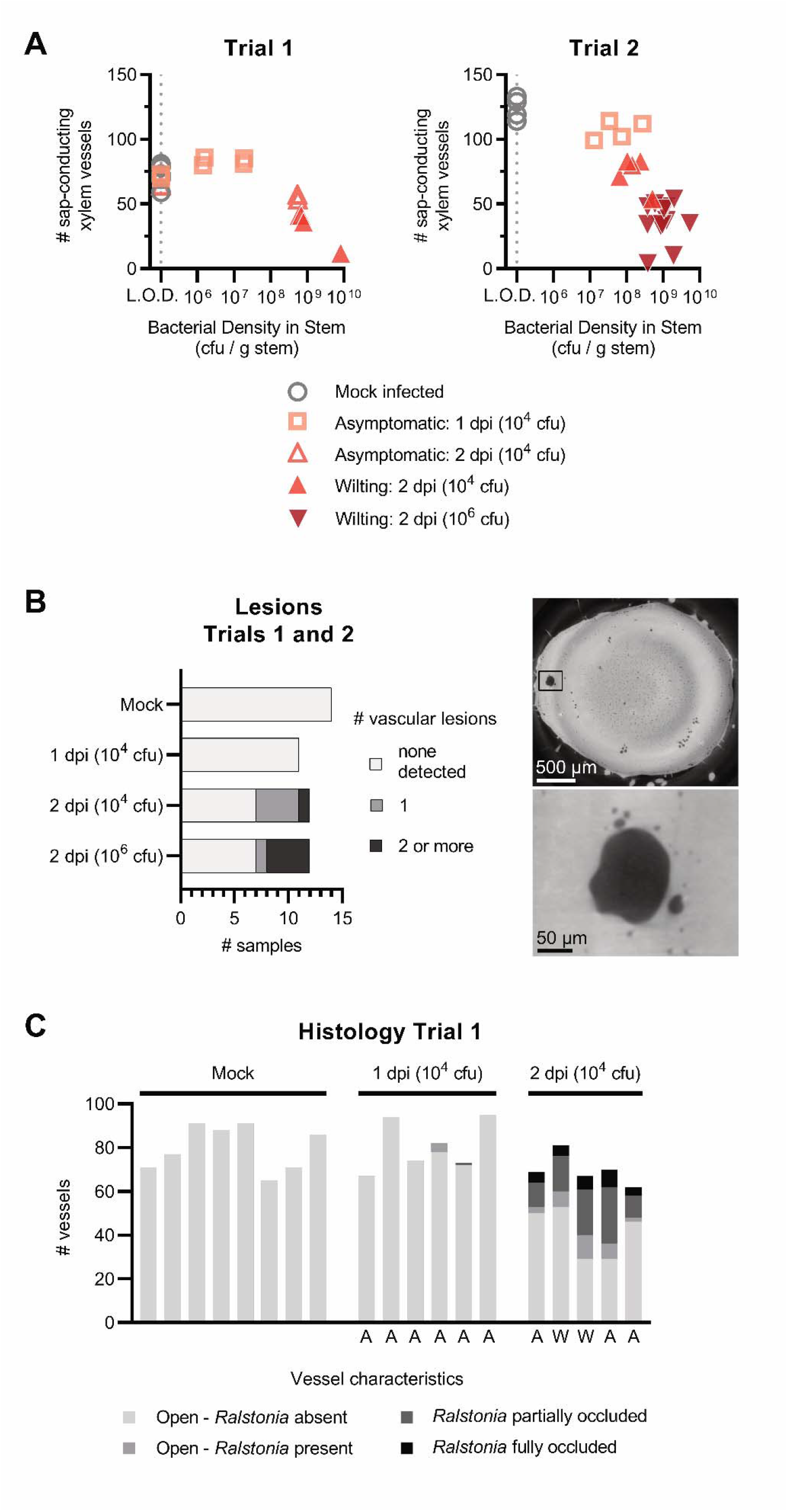
As bacterial wilt disease progresses, fewer vessels are functional, more vessels are occluded by bacterial biomass, and stem lesions develop. (A) The number of sap-conducing xylem vessels and bacterial density from the excised stem imaging assay. The number of sap-conducting (air-filled, black vessels) were quantified from the microCT reconstructions (Fig S5-S8), and bacterial density was quantified by digital droplet PCR. The dashed gray line represents the limit of detection (L.O.D.) of 1×10^5^ CFU/g stem. Samples from mock-inoculated plants were graphed at the L.O.D. (B) Quantification of vessel occlusions in histology micrographs from trial 1 of the excised stem assay. Letters below the graph indicate plant status when stem samples were taken: A = asymptomatic; W = at least one leaf wilting. (C) Vascular lesions near the point of inoculation seen in the reconstructed X-ray microCT cross sections (right and S5-S8) were quantified for each sample in the excised stem assay (left).

Close inspection of the 2 dpi reconstructions revealed that many vessels in the excised stems had dense, white lumina (Fig 3 and S7-S8) similar to those visualized in the stems of intact plants (Fig 2 and S2), indicating that these vessels were occluded. To investigate the occlusions further, we transected the 3D microCT reconstructions to view vessels longitudinally (Fig 5 and S10). When viewed from the side, it was clear that the white occlusions were continuous within the tapered xylem vessels. This suggested that the occlusions were a dense gel formed by bacterial biofilms seen in our and others’ previous microscopic analyses of *Ralstonia*-infected stems (Vasse et al. 1995; Caldwell et al. 2017; Planas-Marquès et al. 2020).

**Fig 5.**
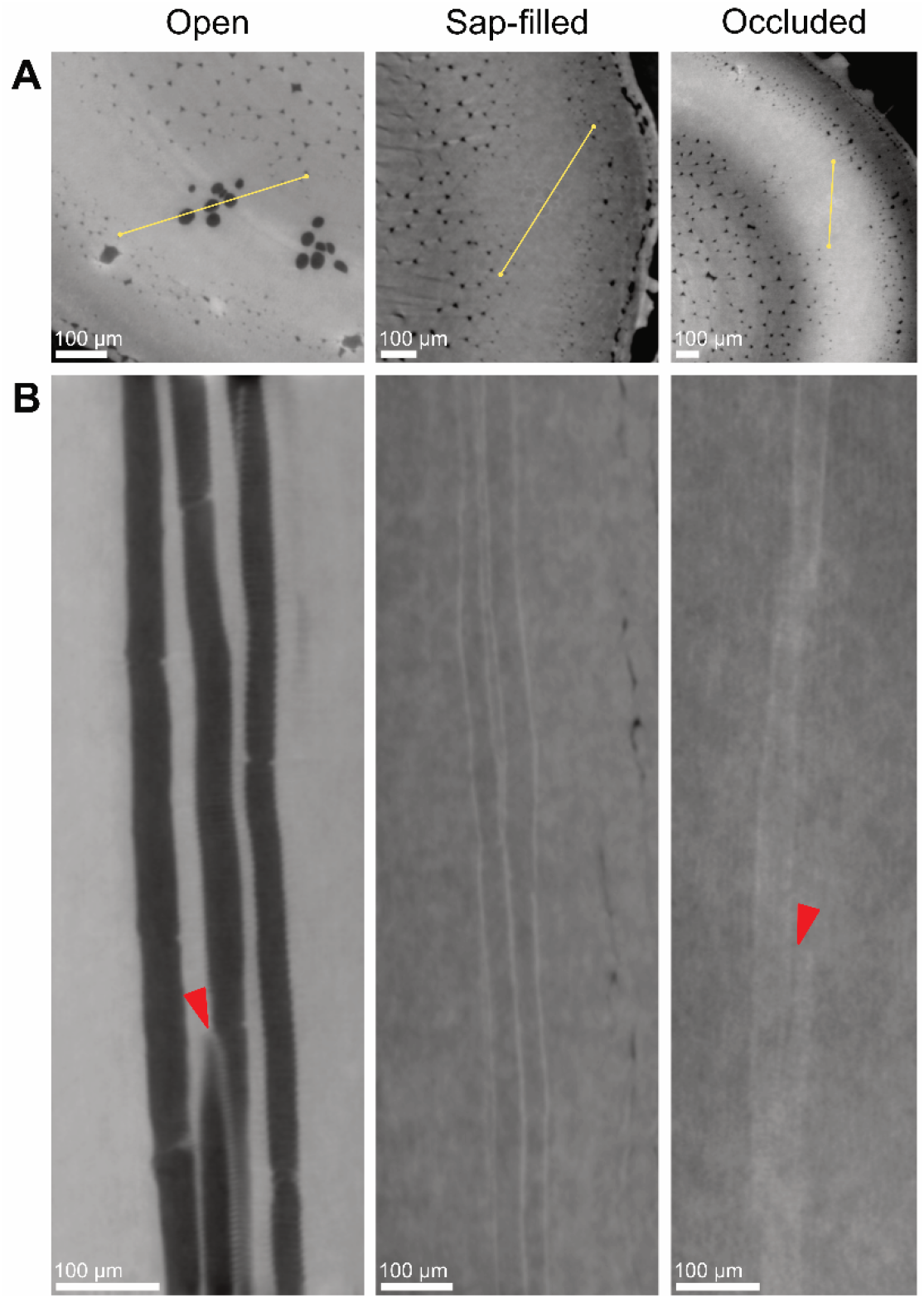
Longitudinal views of xylem vessels. (A) Transverse microCT cross-sections showing open vessels (left), sap-filled vessels (middle), and occluded vessels (right). Yellow lines indicate the axis of longitudinal imaging in B. (B) Stacks of transverse cross sections were compiled and imaged along the longitudinal axes chosen in A. Longitudinal cross-sections of vessels classified as air-filled (left) in transverse cross-sections show that these vessels are filled with air along their length and continuously into connecting vessels via perforation plates (red arrow). Longitudinal cross-sections of vessels classified as sap-filled (middle) in transverse cross-sections show that these vessels are continuously sap-filled along their length. Longitudinal cross-sections of vessels classified as occluded (right) in transverse cross-sections show that these vessels are occluded along their length and continuously into connecting vessels via perforation plates (red arrow). This indicates that the occluding mass is able to move between vessels and clog entire vessel conduits. Samples used in this imaging analysis were H8 (left), HP4 (middle) and 2G4 (right).

Additionally, small lesions were visible near the inoculation site in some of the 2 dpi samples from both trials (Fig 4B), but not in the mock-inoculated or 1 dpi samples. These lesions were irregularly shaped and larger than the 10-40 µm, oval-shaped xylem vessels. In trial 2, these lesions were more common in the 2 dpi plants inoculated with the high titer of bacteria (10^6^ cfu) than the moderate titer (10^4^ cfu). Internal lesions on tomato stems are an infrequent symptom of *Ralstonia* wilt of tomato (García et al. 2019). Lesions like these are often pigmented brown, suggesting that they are part of a plant defense response that involves the activity of polyphenolic oxidases (Beckman 1964).

### Histology of stem samples reveals abundant bacterial occlusions in xylem vessels

We visualized preserved stem samples with a histochemical stain and light microscopy to determine whether the vessel occlusions were plant-derived gels and tyloses or bacterial biofilms. We embedded the stems in paraffin wax and stained slices with 0.05% toluidine blue, which stains the acidic xylem wall cyan and other neutral plant and bacterial tissues dark blue. We classified the lumen contents of each xylem vessel based on the abundance of bacterial biomass within the vessel. Stems from mock-inoculated plants and most 1 dpi plants lacked obvious bacterial aggregates in the xylem (Fig 3D and 4C). By 2 dpi, 23-49% of vessels were partially or fully occluded by *Ralstonia* aggregates, and another 3-16% of xylem vessels were visibly colonized with a few *Ralstonia* cells attached to the walls. Scanning electron microscopy (SEM) analysis of stem tissue confirmed the abundance of bacterial aggregates within xylem vessels (Fig S9). Overall, our results were consistent with the model that the bacterial clogs are the predominant mechanism of vascular disruption during bacterial wilt disease.

## Discussion

Wilt, scorch, blight, and other diseases associated with xylem dysfunction disrupt water transport capacity, causing substantial morbidity and mortality of plants (Beckman et al. 1962; Rajagopal et al. 1987; Denny et al. 1990; Aguirreolea et al. 1995; Lorenzini et al. 1997). Denny et al. (1990) determined that transpiration decreased as wilt symptoms appeared by weighing mock-inoculated and *Ralstonia*-infected tomato plants to measure water loss and estimate whole-plant transpiration rates. In the 1960s, Beckman, Brun, and Buddenhagen constructed an early prototype of a gas exchange system that measured transpiration rates in *Ralstonia*-infected banana plants and demonstrated that *Ralstonia* disrupts water transport (Beckman et al. 1962). We measured the stomatal conductance of individual tomato leaflets to determine transpiration capability on a finer scale instead of a broadly estimating whole plant transpiration. Our results are consistent with the literature that *Ralstonia* wilt symptoms develop only after stomatal conductance decreases. However, stomatal conductance is not solely determined by leaf water potential; chemical signaling via abscisic acid can regulate stomatal aperture independently from leaf water potential (Kim et al. 2010). Therefore, we used X-ray microCT to reveal the ability of individual vessels to transport water.

Although it is well-demonstrated that wilt pathogens disrupt xylem sap transport (Yadeta and Thomma 2013), the exact mechanism(s) of xylem disruption are unknown for most pathosystems. Multiple physiological mechanisms can disrupt xylem water transport: plant-derived structures like tyloses can block vessels, foreign bodies like microbes and their biomass can clog vessels, and gas embolisms can displace sap (Lowe-Power et al. 2018). We used X-ray microCT and complementary methods to test the hypothesis that *Ralstonia* biomass clogs tomato xylem vessels.

Gas embolisms have been associated with drought stress and have been occasionally implicated in early stages of xylem infections (McElrone et al. 2008; Brodersen et al. 2013b; Pérez-Donoso et al. 2016). Embolisms occur when air is aspirated into the water column, which can happen in damaged vascular tissues (Tyree and Sperry 1989). Vascular pathogens often secrete pectinases and cellulases that can degrade the xylem pit membranes that separate vessels from each other and separate vessels from parenchyma tissue (McElrone et al. 2008; Brodersen et al. 2013b; Pérez-Donoso et al. 2016). It has been suggested that some plants are able to refill embolized vessels, though the mechanisms, frequency, and locations of refilling are currently the subject of much debate (Brodersen and McElrone 2013). The phloem-unloading hypothesis proposes that plants could repair embolisms by unloading mono- and disaccharides from the phloem into the xylem (Nardini et al. 2011). These osmolytes would drive water translocation into the vessel, which could dissolve the embolism. If plants divert nutrients into the xylem to ameliorate abiotic stresses, it is likely that some xylem pathogens manipulate this pathway through effectors or by inducing embolisms. We previously found that *Ralstonia-*infection increases the concentration of nutrients in xylem sap (Lowe-Power et al. 2018). We speculated that *Ralstonia’s* degradation of pit membranes (Grimault et al. 1994) might increase vessel embolism, which could trigger an embolism-repair process that increases solute concentration of sap. However, our microCT imaging of intact plants revealed that embolized vessels are rare in mock-infected or *Ralstonia*-infected plants. Clearly, embolism is not responsible for the mass hydraulic failure that induces wilt symptoms, but our data cannot rule out the possibility that vessel embolism occasionally precedes vessel clogging.

We did not observe tyloses in the xylem of *Ralstonia*-infected tomato plants, and our results are consistent with microscopy studies done by multiple groups showing that tyloses are rare in *Ralstonia*-infected tomato plants (Vasse et al. 1995; Rahman et al. 1999; Caldwell et al. 2017). In contrast, MicroCT and microscopy showed abundant Tyloses in xylem vessels of *Xylella-*symptomatic and Esca-symptomatic grapevines (Bortolami et al. 2019; Ingel et al. 2020). Although tyloses can provide an effective defense against some pathogens (Kashyap et al. 2020), prolific tyloses formation in the absence of new vessel development leads to xylem disruption and water stress (Stevenson et al. 2004; Sun et al. 2013; Pérez-Donoso et al. 2016; De Benedictis et al. 2017; Ingel et al. 2020). A preponderance of evidence supports a new model that tyloses, and not bacterial blockages, are responsible for Pierce’s disease symptoms in *Xylella*-infected grapevines (Stevenson et al. 2004; Sun et al. 2013; Pérez-Donoso et al. 2016; De Benedictis et al. 2017; Ingel et al. 2020). We speculate that the differential cause of symptoms between *Ralstonia* and *Xylella* is due to differences in population growth and behavior in their respective host environments. *Ralstonia* can clog xylem vessels by a combination of rapid population growth (exceeding 10^9^ cfu/g within a vessel after a few days) and considerable exopolysaccharide production (McGarvey et al. 1999). Conversely, *Xylella* populations grow at a relatively slow rate (reaching 10^8^ cfu/g after several months) and form aggregates within xylem vessels that often do not occlude the vessel (Baccari and Lindow 2011; Ingel et al. 2020). These factors allow the host plant to mount a defense response within the xylem (e.g. tyloses). However, it has been shown that tyloses are ineffective at containing *Xylella*. Instead, tyloses block an increasing number of vessels over time, resulting in xylem disruption (Ingel et al. 2020). The mechanism of xylem disruption in Esca is still unclear due to the complexity of the disease complex. Esca-diseased grapevines had elevated numbers of tyloses, and experiments with an X-ray contrasting agent in the xylem sap demonstrated that the tyloses were capable of blocking or diverting sap flow. However, the abundance of tyloses did not correlate with symptom severity in the small sample sized study (Bortolami et al. 2019).

Overall, our data support the model that *Ralstonia* infections cause wilt symptoms because the pathogen’s biomass clogs xylem vessels. Importantly, we visualized xylem vessels after both destructive and non-destructive sampling of tomato stems, which allowed us to infer functional status of individual xylem vessels. MicroCT demonstrated that the vessels were clogged in tomato plants with high *Ralstonia* population sizes. Further, histochemical microscopy and SEM showed that bacterial aggregates were abundant in the samples with the most clogged vessels. Consistently, Caldwell et al. (2017) showed histological staining of *Ralstonia*-clogged xylem vessels in susceptible tomato cultivars, while xylem vessels of resistant cultivars remained relatively open. Future studies that incorporate an X-ray contrasting agent like iohexol (Bortolami et al. 2019) with X-ray microCT of *Ralstonia*-infected tomatoes would enable quantification of sap flow rates in xylem vessels with copious vs. minimal *Ralstonia* biomass.

Most of our understanding of plant-microbe interactions is derived from investigating the molecular and cellular level, whole organ, whole plant, or the field level. MicroCT was developed as a medical imaging technology to non-destructively study human physiology and to diagnose human pathologies. MicroCT, MRI, and related technologies allow us to visualize tissues (microCT) and organs (conventional CT) in high resolution 3D. Recently, microCT and magnetic resonance imaging (MRI) has been translated to visualize the physiology and morphology of leafy and woody foliar tissues (Brodersen 2013; Brodersen et al. 2013a; McElrone et al. 2013; Choat et al. 2015; Knipfer et al. 2015; Earles et al. 2018; Li et al. 2019) (Windt et al. 2006). Although vascular plant diseases are the first to be studied with microCT, future studies should visualize the underlying mechanisms of symptom development by foliar pathogens and other root pathogens. Further development and translation of these advanced imaging technologies will enable new insight into plant disease physiology.

## Materials and Methods

### Bacterial growth conditions

Four tomato pathogenic *Ralstonia solanacearum* species complex strains were used in this study: *R. pseudosolanacearum* GMI1000 (phylotype I-18), *R. solanacearum* K60 (IIA-7), *R. solanacearum* UW551 (IIB-1), and *R. solanacearum* UW163 (IIB-4). The bacteria were routinely cultured at 28 °C on CPG media (10 g/L casamino acids, 1 g/L Bacto Peptone, 1 g/L yeast extract, 5 g/L glucose) with 0.002% w/v tetrazolium chloride.

### Measurement of stomatal conductance of tomato leaflets during wilt

Susceptible tomato (*S. lycopersicum* cv. Bonny Best) seed was sown into Sunshine Ready Mix 4 and grown in a growth chamber with constant 28 °C with a 16 hr / 8 hr day / night cycle. After 14 days, seedlings were transplanted into individual 4 inch pots. On day 17, plants were soil-drench inoculated after lightly wounding the roots to synchronize infection (Khokhani et al. 2018). Bacterial inoculum of GMI1000, UW551, K60, or UW163 was prepared from overnight cultures; cells were pelleted by centrifugation and resuspended in water to OD = 0.2. Roots were lightly wounded by gently lifting the plant stem, and 50 ml of the inoculum was poured into the soil. Control plants were wounded and inoculated with pure water.

At 3 - 4 days post inoculation, stomatal conductance was measured on individual leaflets of partially-wilted *Ralstonia*-infected plants and on mock-inoculated non-wilted plants with a Licor 6400XT gas analyzer outfitted with a broadleaf chamber. Measurements occurred over the course of 3.5 hr at 2-5.5 post-light-onset within the growth chamber. The Licor 6400XT was run with settings that matched the measured environmental values of the growth chamber: 700 ppm reference CO_2_, 350 µmol m^-2^ s^-1^ light intensity, 400 μmol s^-1^ airflow, chamber temperature at 27 °C to match thermocouple measurements of leaf temperature. Individual leaflets were enclosed in the gas analyzer chamber and stomatal conductance was recorded at equilibrium. To account for diurnal changes in stomatal conductance, mock-inoculated and *Ralstonia*-infected plants were measured in alternating blocks of 1-2 plants. On each mock-inoculated plant, 3-4 leaflets on individual petioles were measured; all leaflets of these plants were turgid. Leaflets on *Ralstonia*-infected plants with three phenotypes were targeted for measurements: “turgid” leaflets located on a petiole without any wilted leaflets, “wilted” leaflets, and “turgid near wilted” which were visually turgid leaflets that shared a petiole with wilted leaflets (Fig 1). The sample size per condition varied based on wilt symptom distribution the plant, *i*.*e*. not all symptomatic plants had petioles with both wilted and turgid leaflets.

### X-ray Microcomputed Tomography

To prepare infected plants with a range of bacterial burdens and mock-inoculated plants to be imaged by X-ray microcomputed tomography during a single 8-12 hr imaging session, plants were inoculated by the cut-petiole method 1 or 2 days before the imaging session (Khokhani et al. 2018). This cut-petiole inoculation method was used because it yields highly synchronized disease progression relative to the stochastic disease progression of soil-drench inoculations. Strain GMI1000 was chosen for these studies because it is a model *R. pseudosolanacearum* strain.

Susceptible tomato (*S. lycopersicum* cv. Money Maker) seed was sown into Sunshine Ready Mix 4 and grown in a greenhouse. After 14 days, seedlings were transplanted into individual 4 inch pots and returned to the greenhouse. At 25 and 26 days post sowing, tomato plants with 3-4 petioles were inoculated by excising the oldest petiole with a razor blade and carefully reverse-pipetting a 2 μl bacterial suspension (1 x 10^4^ or 1×10^6^ total colony forming units per plant). Mock inoculated plants were inoculated with a droplet of sterile water. Inoculated plants were then grown for 1-2 days in an E15 Conviron growth chamber with constant 28 °C and 16 / 8 hr day/light cycle.

Plants were imaged at the Lawrence Berkeley National Laboratory Advanced Light Source (ALS) microCT facility (beamline 8.3.2). Samples were continuously rotated 180° while scanned in a 21 keV synchrotron X-ray beam as described in McElrone et al. (2013). Per 180° rotation, 769-1313 longitudinal images were acquired by a CMOS camera (PCO.edge; PCO AG, Kehlheim, Germany) at 250 -500 ms exposure time. Image resolution was 1.3 μm or 3.25 μm per pixel depending on whether the stem diameter allowed it to be scanned at 5x or 2x magnification. Reconstructed cross-section images were analyzed in ImageJ.

Two experimental designs were used to visualize anatomy and physiology of the stem vasculature during bacterial wilt disease.

### Experiment 1: Visualization of vascular system of plants with intact roots and leaves

Intact plants at the 3 leaf stage were stabilized between Styrofoam blocks on a custom rig to prevent movement during imaging. The stem was imaged 2 mm above the inoculation site. 3 mock inoculated and 7 *R. pseudosolanacearum* GMI1000 inoculated plants with a range of symptoms were imaged. Reconstructed cross-section images were analyzed in ImageJ to quantify the number of embolized vessels (appear black in the images) (Fig S11A).

### Experiment 2: Visualization of vascular system of dehydrated, excised stem

To visualize the water-conducting capacity of vessels, a 2.5 cm stem segment was cut 1.25 cm above and below the inoculated petiole which exposed the disrupted xylem vessels to air. The segments were briefly dehydrated under a heat lamp for 25 min, which allowed the sap-conducting vessels to empty out. In contrast, clogged vessels would not empty. The stem segments were then loaded onto a drill chuck microCT rig and imaged. Reconstructed cross-section images were analyzed in ImageJ to determine the number of unclogged vessels (appear black in the images) and clogged vessels (appear whiter than the background tissue in the images) (Fig S11B).

### Bacterial quantification by digital droplet PCR

A *Pseudomonas syringae* digital droplet PCR (ddPCR) protocol (Morella et al. 2018) was modified to quantify *Ralstonia solanacearum* population sizes in tomato stem homogenate. Universal *Ralstonia* primers 759 (5’-GTC GCC GTC AAC TCA TT TCC) and 760 (5’-GTC GCC GTC AGC AAT GCG GAA TCG) that amplify a 281 bp band (Opina et al. 1997). A mastermix was prepared with of QX200 ddPCR EvaGreen Supermix (Bio-Rad), 250 nM each of 759/760 primers, and 2 ul of template (pure DNA or bacterial cells for a colony PCR approach). Oil-encapsulated droplets were generated in a Bio-Rad droplet generator per manufacturer’s protocol before PCR amplification. Thermocycler conditions for the PCR were 95 °C 2 min; 40 cycles: 94 °C 20 s, 60 °C 20s, 72 °C 15 s; 72 °C 5 min. Amplified samples were analyzed on the Bio-Rad droplet reader following manufacturer’s instructions and Bio-Rad software was used for data analysis as described (Morella et al. 2018).

Before quantifying bacterial populations in the microCT imaged experimental samples, the ddPCR protocol was optimized as follows. Briefly, standard curves of *R. solanacearum* cells at known concentrations (1×10^3^ to 1×10^5^ cells per ddPCR reaction) were spiked into tomato stem homogenate (30 mg stem in 1 ml water), a 1:10 dilution of the homogenate, or water. The undiluted homogenate inhibited the PCR reaction, but reactions of the 1:10 homogenate and water accurately reflected bacterial concentration.

For experimental samples, stem tissue (19-40 mg) just above the site of inoculation was placed on wet ice immediately after microCT imaging and were frozen within 12 hrs. Samples were thawed and homogenized in 1 ml water in a BioSpec Mini BeadBeater-8 shaking bead mill with four 2.2 mm metal beads for 90s. Homogenate was statically incubated for 5 min to allow plant biomass to settle before diluting the sample 1:10. Two μl of the diluted homogenate was used as template DNA in the reactions.

### Microscopy & counting vessels

Samples from one trial of the excised stem experiment were preserved for histochemical staining and light microscopy. After imaging, the portion of the stem tissue that was secured in the drill chuck was preserved in 70% ethanol. Samples were stored at room temperature until prepared for microscopy as described in Caldwell et al. (2017). Briefly, samples were further dehydrated with a graded series of alcohols, embedded in paraffin, and further mounted in the microtome sample holder. Once the paraffin block was constructed, the sample was cut using a rotary microtome and the cut samples were placed on a glass slide for light microscopy. A polychromatic dye, 0.05% Toluidine Blue, was used to assist in visualization of bacteria within plant tissues. When a region of interest was identified, paraffin samples were then recut using protocol described in Caldwell et al. 2019. Light microscopy images were captured using an Olympus CX43 microscope and the SPOT Idea CMOS Camera. Images were sent to UC Davis for analysis and counting. Vessels were visually inspected for the presence of *Ralstonia* and counted based on the degree of *Ralstonia* colonization: Open-absent – an open vessel without *Ralstonia*; Open-*Ralstonia* present – an open vessel with a low concentration of *Ralstonia* congregating at the vessel wall; *Ralstonia* partially occluded – a vessel where *Ralstonia* has colonized beyond the vessel wall and is forming an occlusion, but has not fully occluded the vessel; *Ralstonia* fully occluded – a vessel that is fully occluded by *Ralstonia*.

## Data availability

Representative, high resolution cross-sections of each microCT reconstruction are available at 10.6084/m9.figshare.13287290 and 10.6084/m9.figshare.13283933. The tomopy832.py script used to reconstruct the X-ray data is available https://gist.github.com/lbluque/1bed77425fc531a882cfcca68dd3cb6e. The raw X-ray scatter data (1-10 gigabyte per file) and input parameters used in the reconstructions are available upon request.

## Acknowledgements

We sincerely thank multiple scientists for technical advice: Duncan Smith for Licor 6400XT training, Thorsten Knipfer, Clarissa Reyes, Alastair MacDowell, and Harold Barnard for microCT training and support, and Jason Lowe-Power for computational training. Dr. Caitilyn Allen provided the facilities to measure stomatal conductance for all strains, including the R3B2 select agent strain UW551. Dr. Steven Lindow and Christina Wistrom provided access to and support for CNR Oxford Tract Greenhouse where plants were grown for X-ray analysis.

Funding was provided to T. Lowe-Power (USDA NIFA Postdoctoral fellowship #15148) and the Iyer-Pascuzzi laboratory (FFAR New Innovator Award). This research used resources of the Advanced Light Source, which is a DOE Office of Science User Facility under contract no. DE-AC02-05CH11231.

---

**Fig S1.**
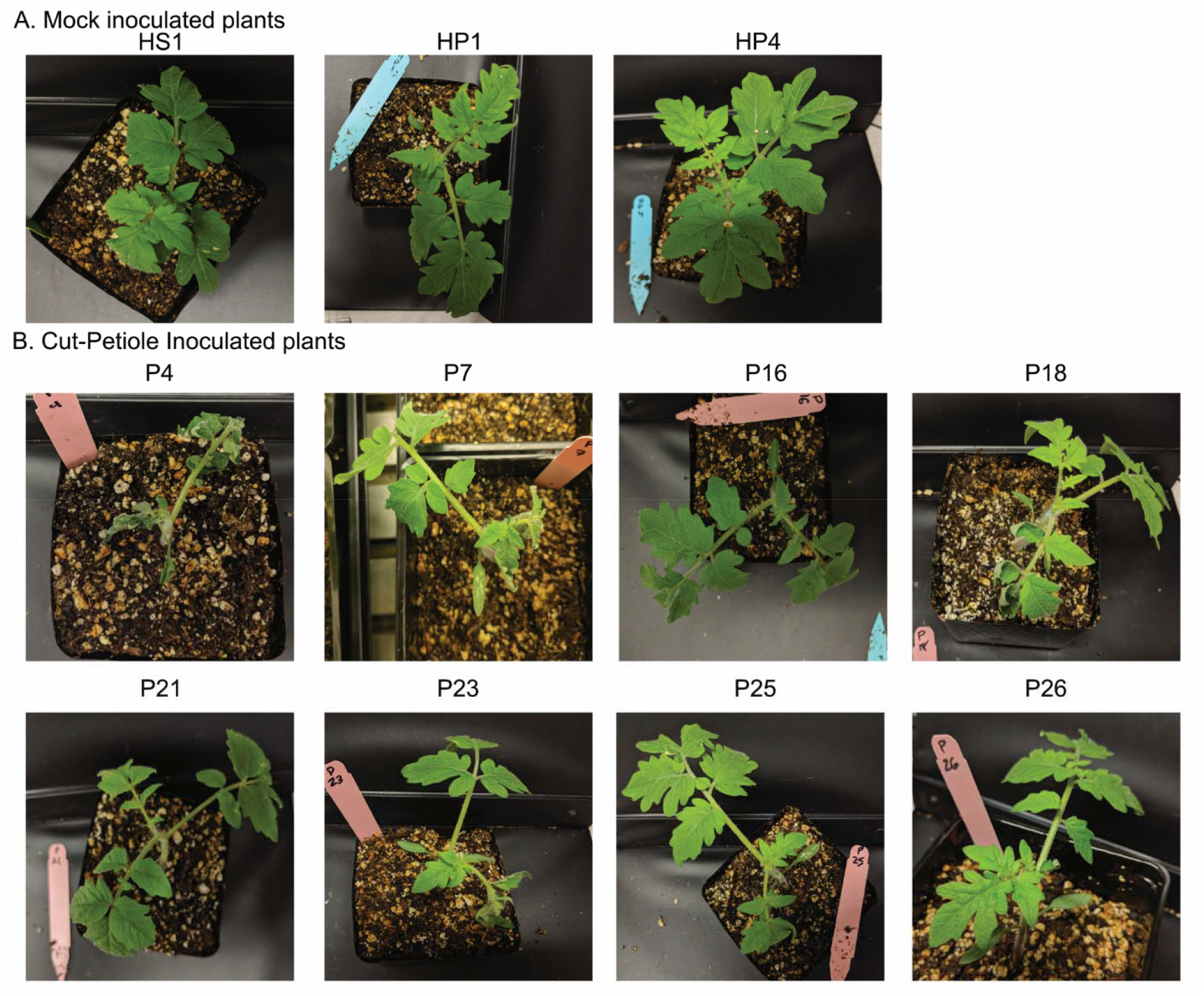
Photographs of plants imaged as intact plants. Photographs of mock-inoculated and *Ralstonia*-infected plants before microCT imaging. These images correspond to representative data in Fig 2 and the full set of microCT images in Fig S2. Plants were assigned alphanumeric identifiers to match images between supplemental figures.

**Fig S2.**
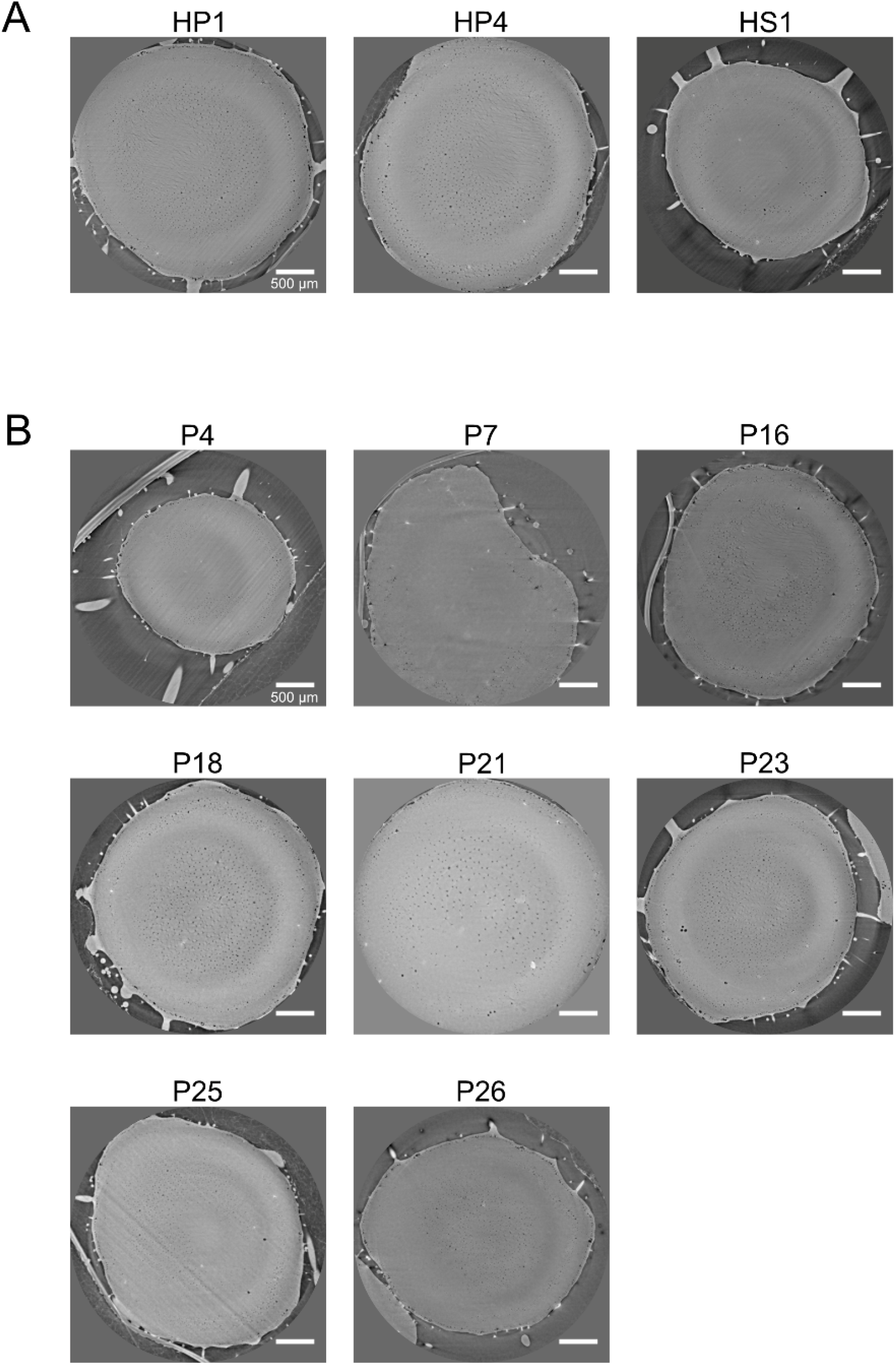
MicroCT Cross-sections of stems from intact plants. MicroCT cross-sections correspond to representative data in Fig 2 and whole plant photographs in Fig S1. Plants were assigned alphanumeric identifiers to match images between supplemental figures.

**Fig S3.**
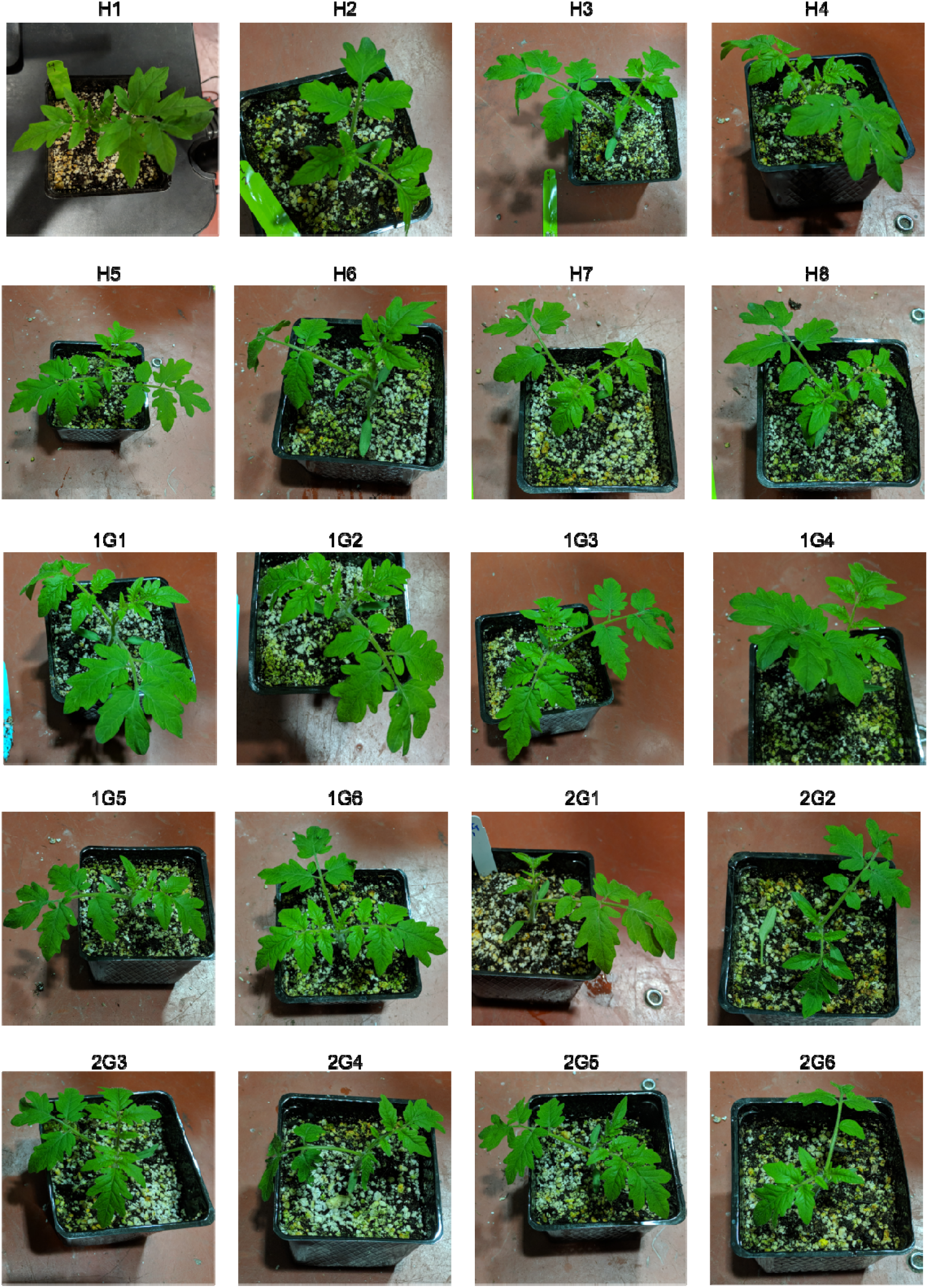
Photographs of whole plants used in trial 1 of the excised stem assay. Photographs of mock-inoculated and *Ralstonia*-infected plants before microCT imaging of excised stems. Photographs of mock-inoculated plants and plants inoculated with 10^4^ cfu *Ralstonia* were taken 1-2 days before microCT imaging. These photographs correspond to representative data in Fig 3 and microCT images in Fig S5-S8. Plants were assigned alphanumeric identifiers to match images between supplemental figures. “H-” indicates mock-inoculated plants, “1G-” indicates *Ralstonia-*infected plant 1 dpi with 10^4^ cfu, and “2G-” indicates *Ralstonia-*infected plant 2 dpi with 10^4^ cfu.

**Fig S4.**
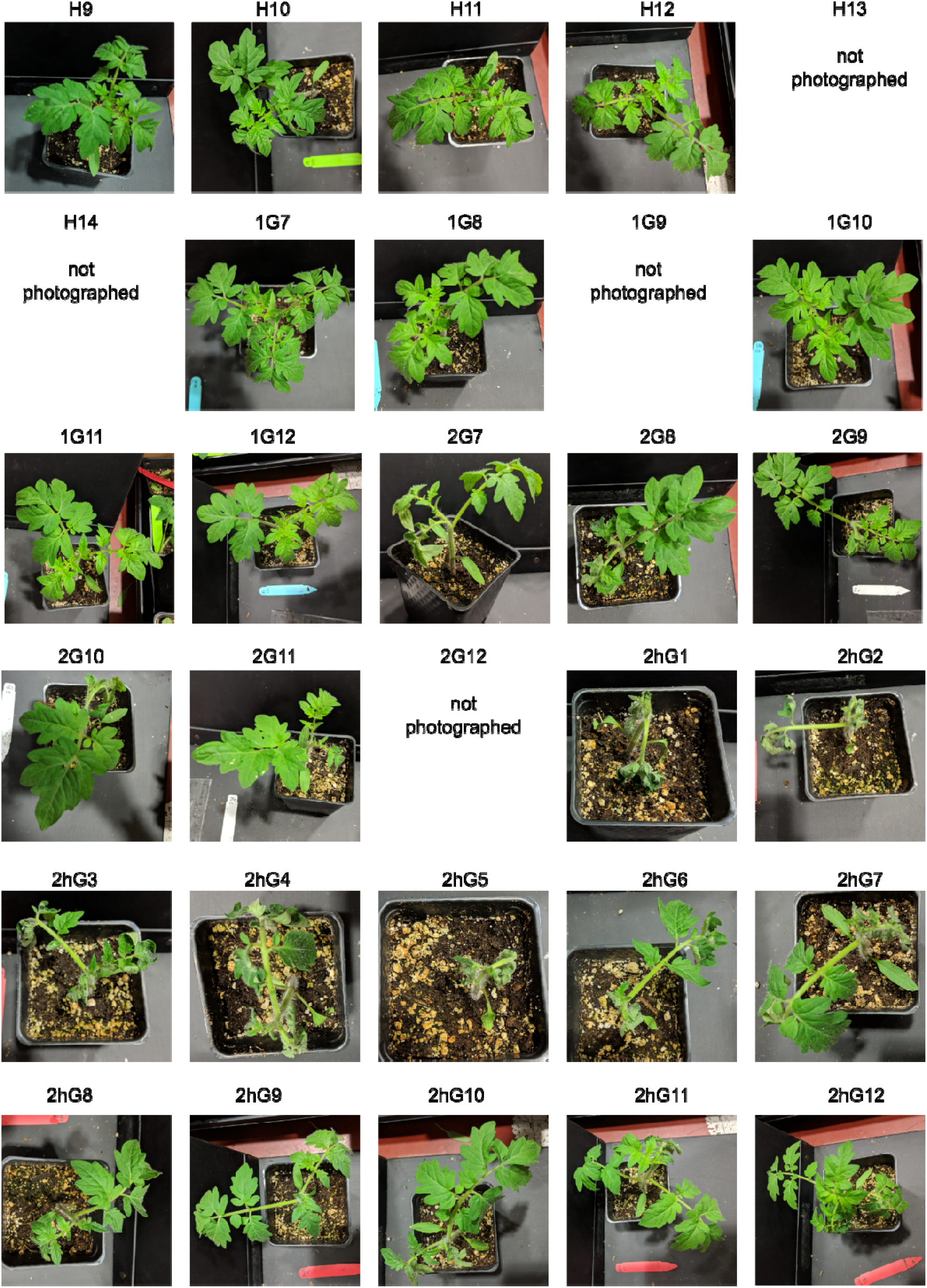
Photographs of whole plants used in trial 2 of the excised stem assay. Photographs of mock-inoculated and *Ralstonia*-infected plants before microCT imaging of excised stems. Photographs of mock-inoculated plants and plants inoculated with 10^4^ or 10^6^ cfu *Ralstonia* were taken 1-2 days before microCT imaging. These images correspond to representative data in Fig 3 and microCT images in Fig S5-S8. Plants were assigned alphanumeric identifiers to match images between supplemental figures. “H-” indicates mock-inoculated plants, “1G-” indicates *Ralstonia-*infected plant 1 dpi with 10^4^ cfu, “2G-” indicates *Ralstonia-*infected plant 2 dpi with 10^4^ cfu, and “2hG-” indicates *Ralstonia-*infected plant 2 dpi with the higher 10^6^ cfu titer. Four plants were “not photographed” before the stem was destructively excised.

**Fig S5.**
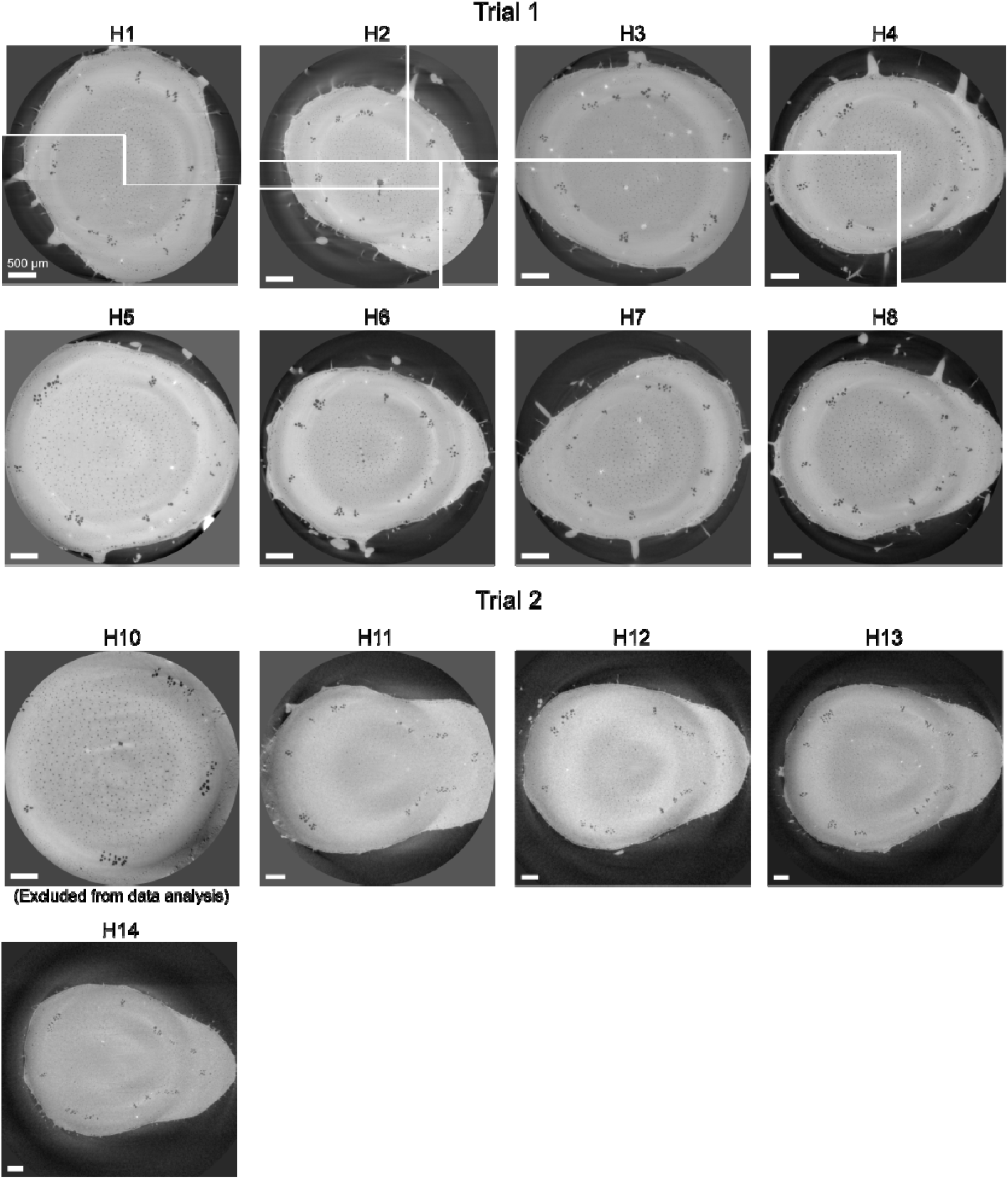
MicroCT cross-sections of mock-inoculated plant stems after excision and dehydration. These images correspond to representative data in Fig 3 and whole plant images in Fig S3-S4. Plants were assigned alphanumeric identifiers to match images between supplemental figures. “H-” indicates mock-inoculated plants. Plants were analyzed in 2 trials with 8 mock-inoculated plants in trial 1 and 6 plants in trial 2. Initial scans (H1 to H4) were higher resolution (more images acquired per 180° rotation) and lengthy imaging time allowed the stem to further dehydrate. This dehydration shrinkage shifted the center of rotation (cor), so multiple cor inputs were required to focus around the cross-section. Different reconstruction parameters are indicated by gaps in the image. Sample H9 was imaged out-of-frame and not reconstructed. Samples with the xylem outside of the frame of view (H10) were not included in quantitative analyses. Scale bars indicate 500 μm.

**Fig S6.**
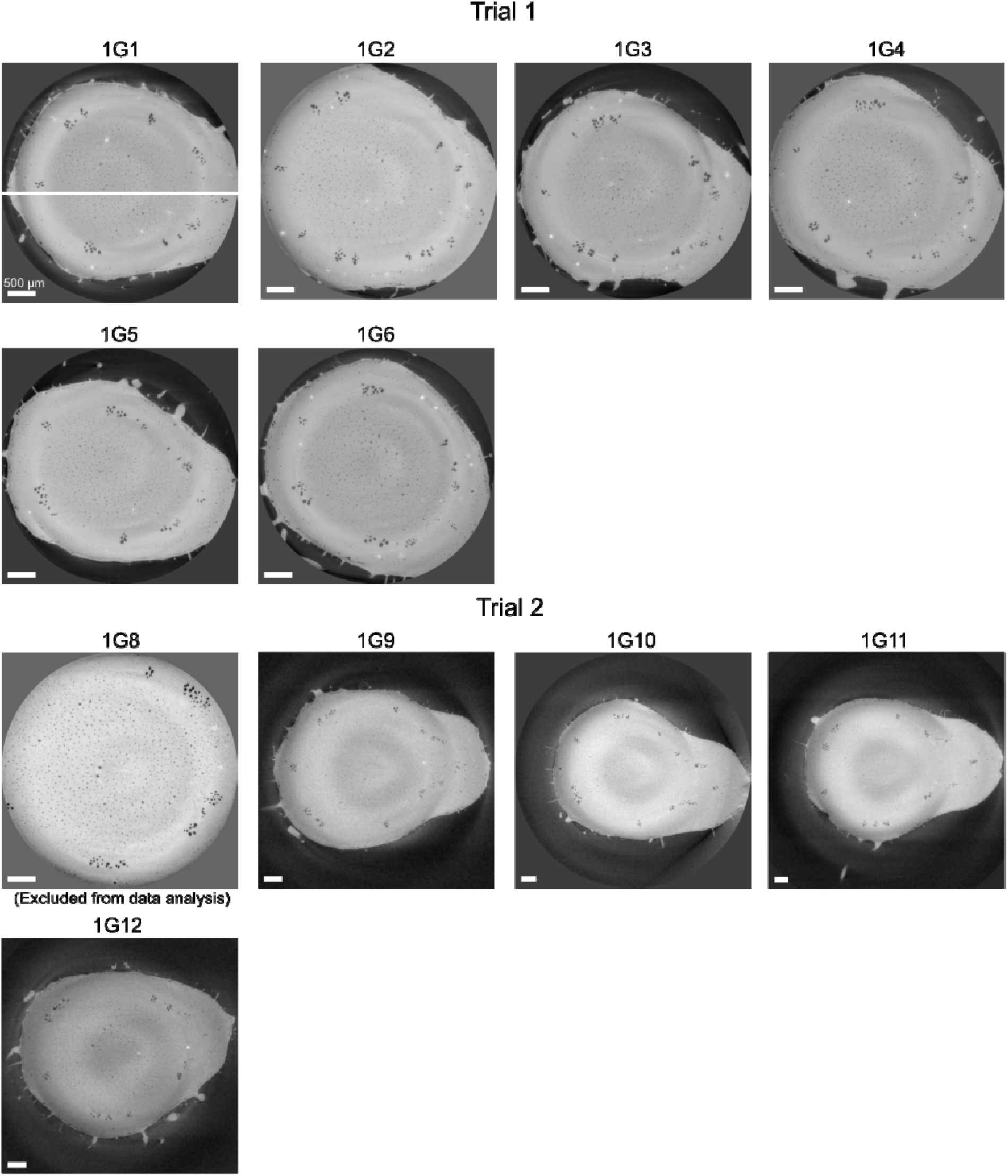
MicroCT cross-sections of 1 dpi infected plant stems after excision and dehydration. These images correspond to representative data in Fig 3 and whole plant images in Fig S3-S4. Plants were assigned alphanumeric identifiers to match images between supplemental figures. “1G-” indicates *Ralstonia-*infected plant 1 dpi with 10^4^ cfu inoculum. Plants were analyzed in 2 trials with 6 plants per trial. Initial scans (1G1) were higher resolution (more images acquired per 180 rotation), and lengthy imaging time allowed the stem to further dehydrate. This dehydration shrinkage shifted the center of rotation (cor), so multiple cor inputs were required to focus around the cross-section. Different reconstruction parameters are indicated by gaps in the image. Sample 1G7 was out-of-focus and not reconstructed or included in any analysis. Samples with the xylem outside of the frame of view (1G8) were not included in quantitative analyses. Scale bars indicate 500 μm.

**Fig S7.**
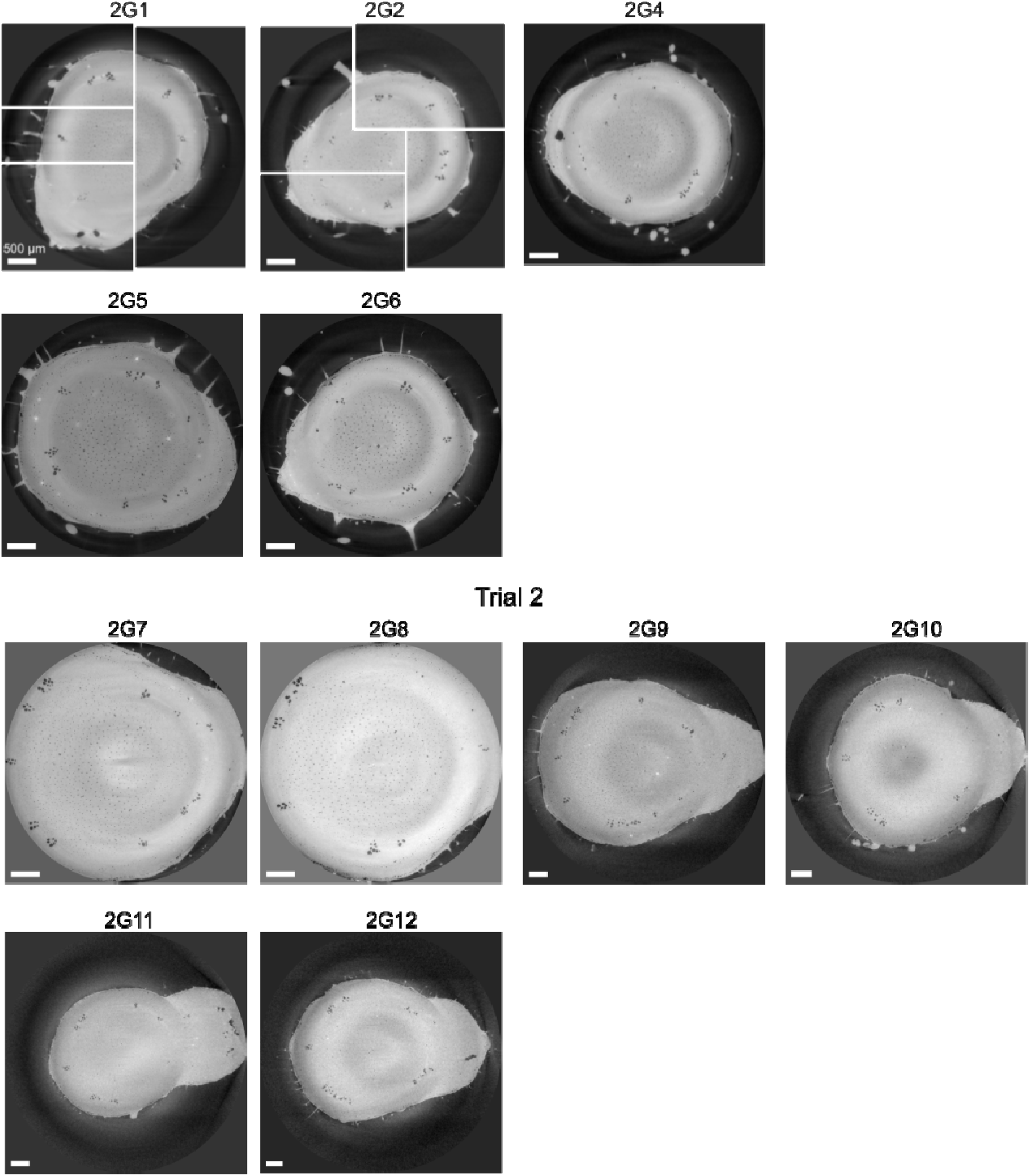
MicroCT cross-sections of 2 dpi infected plant stems after excision and dehydration (10^4^ cfu inoculum) These images correspond to representative data in Fig 3 and whole plant images in Fig S3-S4. Plants were assigned alphanumeric identifiers to match images between supplemental figures. “2G-” indicates Ralstonia-infected plant 2 dpi with 10^4^ cfu inoculum. Plants were analyzed in 2 trials with 5-6 plants each. Initial scans (2G1 to 2G2) were higher resolution (more images acquired per 180 rotation), and lengthy imaging time allowed the stem to further dehydrate. This dehydration shrinkage shifted the center of rotation (cor), so multiple cor inputs were required to focus around the cross-section. Different reconstruction parameters are indicated by gaps in the image. Scale bars indicate 500 μm.

**Fig S8.**
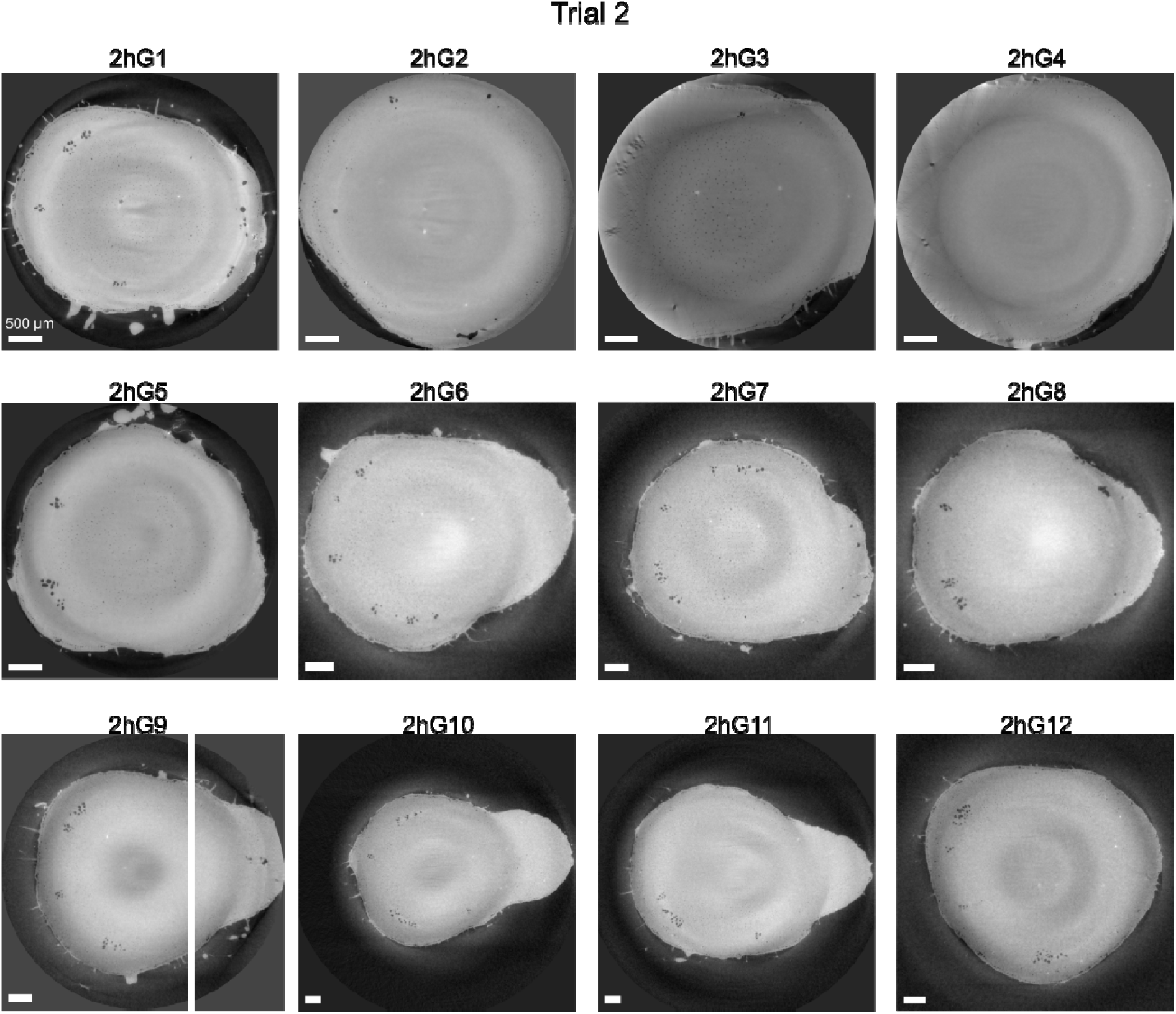
MicroCT cross-sections of 2 dpi infected plant stems after excision and dehydration (10^6^ cfu inoculum) These images correspond to representative data in Fig 3 and whole plant images in Fig S3-S4. Plants were assigned alphanumeric identifiers to match images between supplemental figures. “2hG-” indicates Ralstonia-infected plant 2 dpi with the higher 10^6^ cfu inoculum. Plants were analyzed in as one trial with 12 plants (imaged as part of trial 2). Sample 2hG3 and 2hG4 were rotated near the periphery of the detector and cannot be perfectly reconstructed. Nevertheless, air-filled xylem vessels were countable despite crescent-shaped distortions, so 2hG3 and 2hG4 were included in quantitative analyses. Sample 2hG9 required two center of rotation (cor) inputs and a composite image is displayed. Scale bars indicate 500 μm.

**Fig S9.**
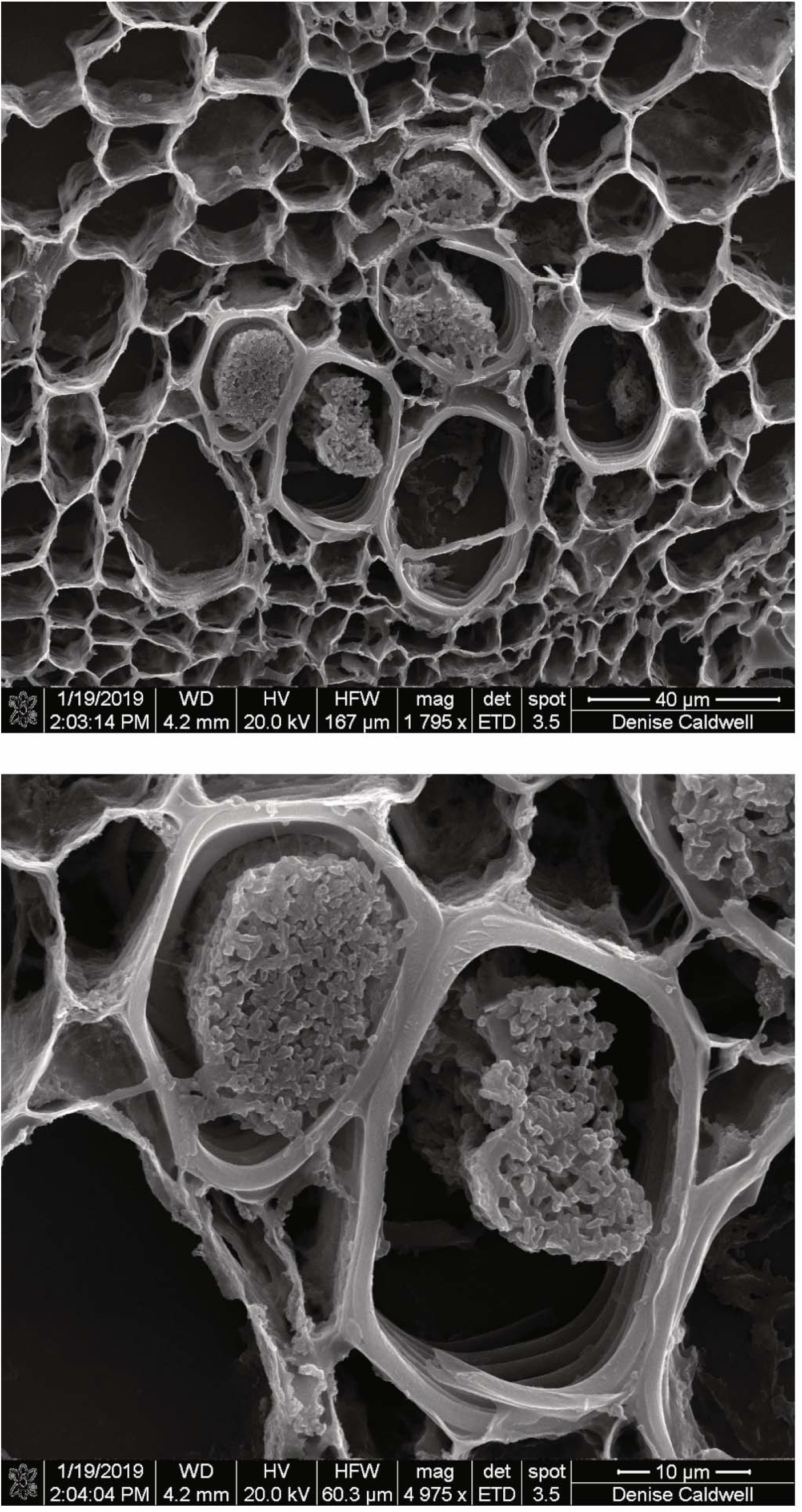
Scanning electron microscopy of 2 dpi infected plant stems after excision and dehydration (10^4^ cfu inoculum). Scanning electron microscopy (SEM) of an excised stem from a 2 dpi infected plant (2G2) shows tomato xylem vessels that appear to be partially or fully occluded by *Ralstonia*. The spatial arrangement of partial and fully occluded vessels captured by SEM is consistent with that visualized by microCT and light microscopy with histological staining.

**Fig S10.**
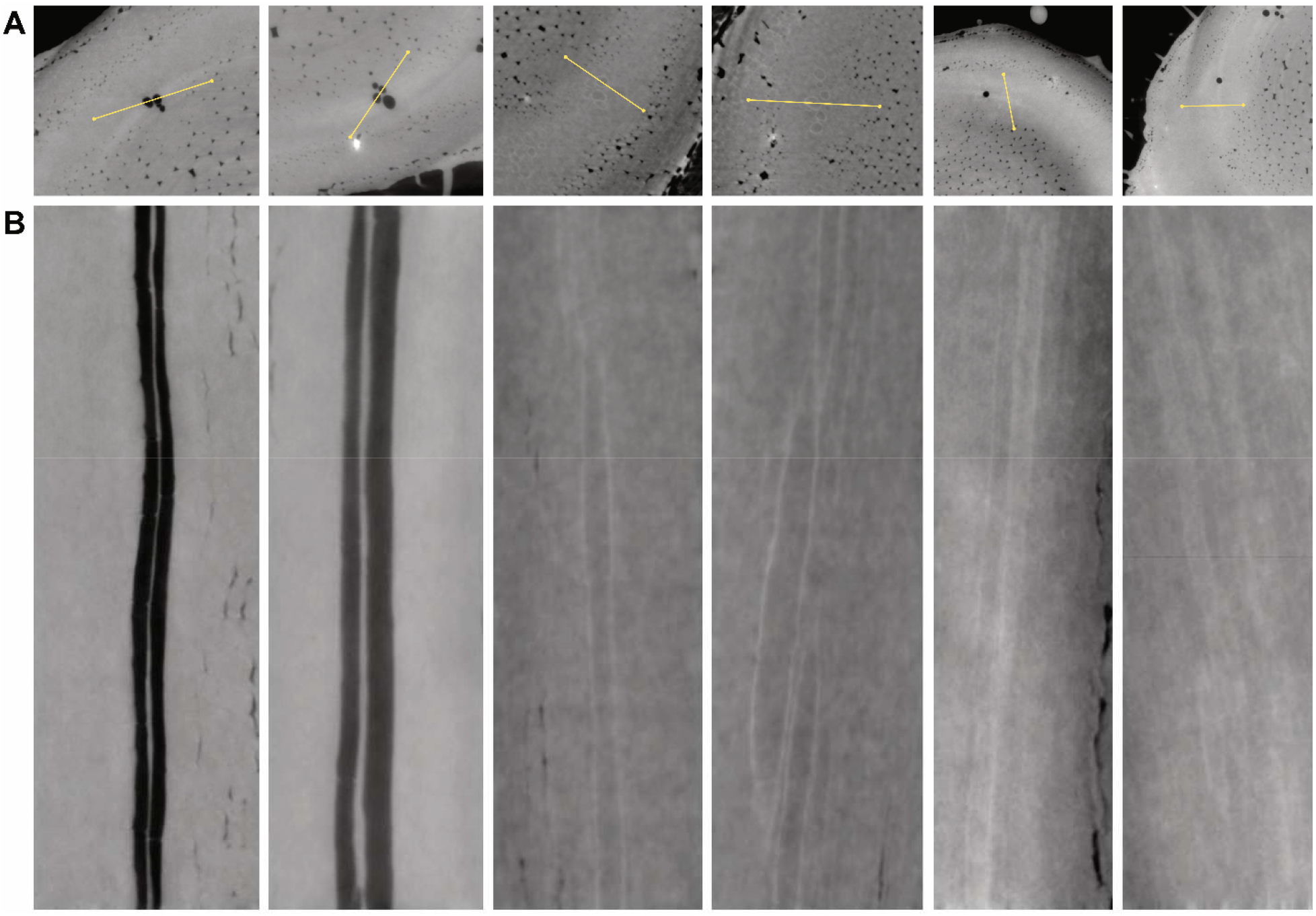
MicroCT cross-sections and corresponding longitudinal sections. (A) cross sections of samples (left to right): H7, H8, HP4, HP4, 2G4, 2G6. Yellow lines indicate the axis of imaging for longitudinal sections. (B) Longitudinal sections depicting open, empty vessels (H7, H8), sap-filled vessels (HP4) and vessels with dense occlusions (2G4, 2G6).

**Fig S11.**
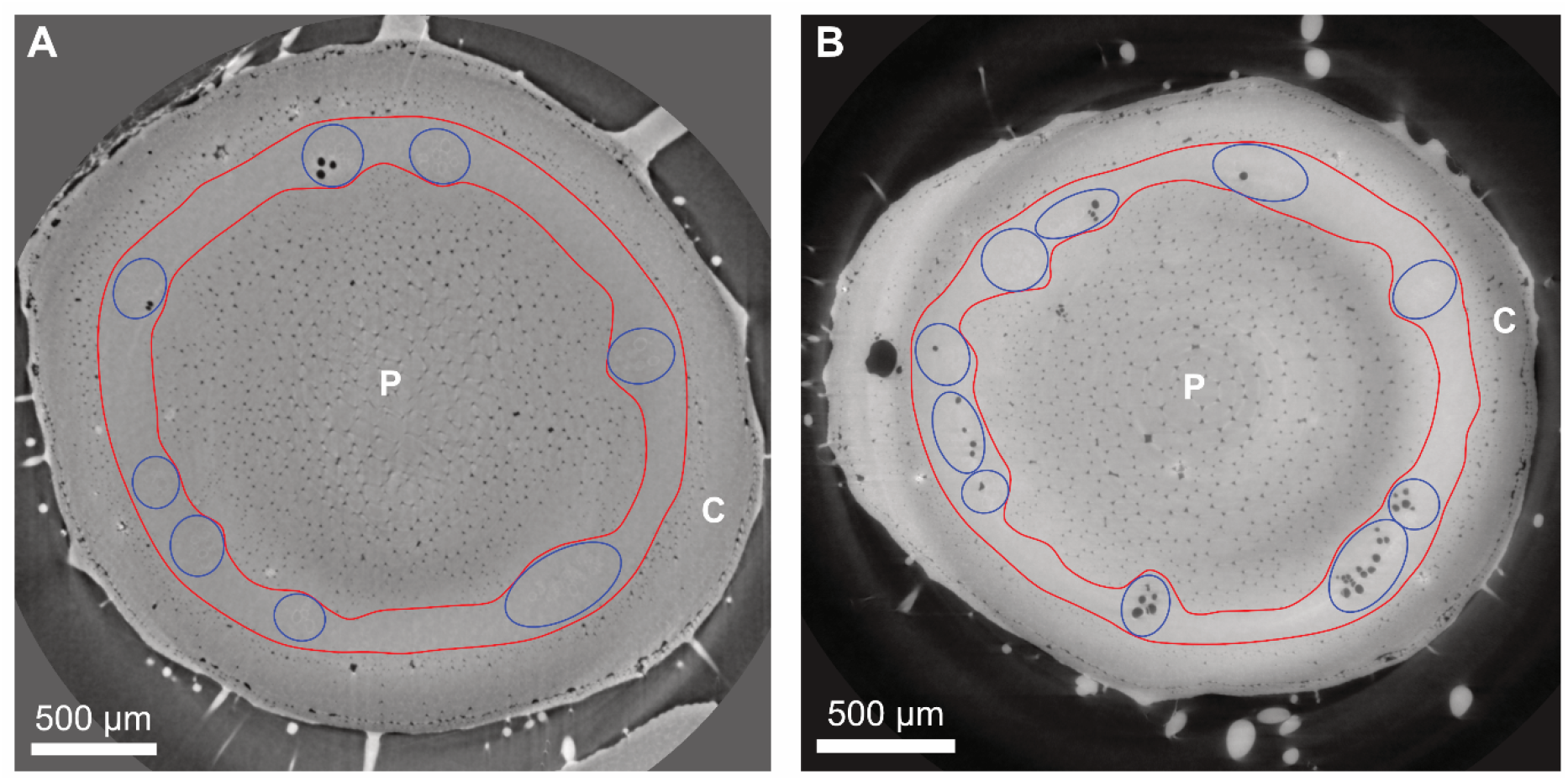
Methodology for counting tomato xylem vessels in microCT images. The vasculature where xylem vessels are located is shown between the two red lines for (A) intact stem microCT images, and (B) excised stem microCT images. The area in the middle is the pith (P) and the area outside the vasculature is the cortex (C). Xylem vessel bundles are indicated by blue ellipses and individual vessels were counted from these bundles.

**Fig S12.**
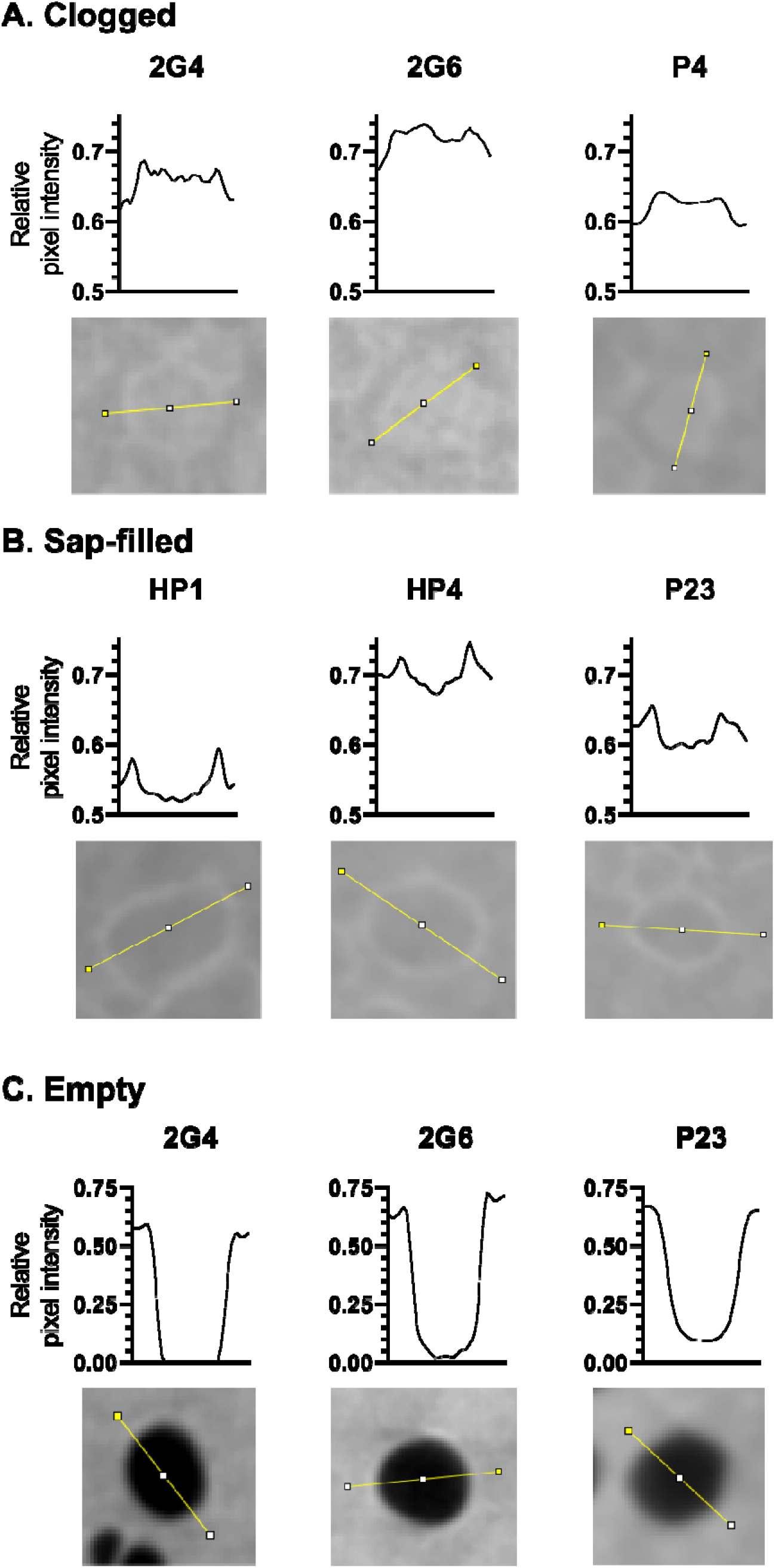
Relative pixel intensity of clogged, sap-filled, and empty xylem vessels. The relative pixel intensity of xylem vessels was calculated using ImageJ from microCT images along a linear axis (yellow line) for clogged vessels (A), sap-filled vessels (B), and empty vessels (C), where zero = black and 1 = white. (A) Clogged vessels displayed higher pixel intensities relative to the surrounding tissue and had pixel intensities similar to dense xylem vessel walls. (B) Sap-filled vessels displayed similar pixel intensities to the surrounding tissue and contrasted with dense vessel walls that had higher pixel intensity. (C) Gas-filled vessels displayed pixel intensities near zero.

## References

Aguirreolea, J., Irigoyen, J., Sánchez-Díaz, M., and Salaverri, J. 1995. Physiological alterations in pepper during wilt induced by Phytophthora capsici and soil water deficit. Plant Pathol. 44:587–596

Baccari, C., and Lindow, S. E. 2011. Assessment of the process of movement of Xylella fastidiosa within susceptible and resistant grape cultivars. Phytopathology. 101:77–84

Beckman, C. H. 1964. Host responses to vascular infection. Annu. Rev. Phytopathol. 2:231–252

Beckman, C. H., Brun, W. A., and Buddenhagen, I. W. 1962. Water Relations in Banana Plants Infected with Pseudomonas solanacearum. Phytopathology. 52:1144

Bortolami, G., Gambetta, G. A., Delzon, S., Lamarque, L. J., Pouzoulet, J., Badel, E., Burlett, R., Charrier, G., Cochard, H., Dayer, S., Jansen, S., King, A., Lecomte, P., Lens, F., Torres-Ruiz, J. M., and Delmas, C. E. L. 2019. Exploring the Hydraulic Failure Hypothesis of Esca Leaf Symptom Formation. Plant Physiol. 181:1163–1174

Brodersen, C. R. 2013. Visualizing wood anatomy in three dimensions with high-resolution X-ray micro-tomography (μCT) – A review. Pages 80–96 in: Wood Structure in Plant Biology and Ecology, Brill.

Brodersen, C. R., Choat, B., Chatelet, D. S., Shackel, K. A., Matthews, M. A., and McElrone, A. J. 2013a. Xylem vessel relays contribute to radial connectivity in grapevine stems (Vitis vinifera and V. arizonica; Vitaceae). Am. J. Bot. 100:314–321

Brodersen, C. R., and McElrone, A. J. 2013. Maintenance of xylem Network Transport Capacity: A Review of Embolism Repair in Vascular Plants. Front. Plant Sci. 4:108

Brodersen, C. R., McElrone, A. J., Choat, B., Lee, E. F., Shackel, K. A., and Matthews, M. A. 2013b. In vivo visualizations of drought-induced embolism spread in Vitis vinifera. Plant Physiol. 161:1820–1829

Brodribb, T. J., McAdam, S. A., and Carins Murphy, M. R. 2017. Xylem and stomata, coordinated through time and space. Plant Cell Environ. 40:872–880

Caldwell, D., Kim, B.-S., and Iyer-Pascuzzi, A. S. 2017. Ralstonia solanacearum Differentially Colonizes Roots of Resistant and Susceptible Tomato Plants. Phytopathology. 107:528–536

Choat, B., Brodersen, C. R., and McElrone, A. J. 2015. Synchrotron X-ray microtomography of xylem embolism in Sequoia sempervirens saplings during cycles of drought and recovery. New Phytol. 205:1095–1105

Choat, B., Cobb, A. R., and Jansen, S. 2008. Structure and function of bordered pits: new discoveries and impacts on whole-plant hydraulic function. New Phytol. 177:608–625

Dalsing, B. L., and Allen, C. 2014. Nitrate assimilation contributes to Ralstonia solanacearum root attachment, stem colonization, and virulence. J. Bacteriol. 196:949–960

Dalsing, B. L., Truchon, A. N., Gonzalez-Orta, E. T., Milling, A. S., and Allen, C. 2015. Ralstonia solanacearum uses inorganic nitrogen metabolism for virulence, ATP production, and detoxification in the oxygen-limited host xylem environment. MBio. 6:e02471

De Benedictis, M., De Caroli, M., Baccelli, I., Marchi, G., Bleve, G., Gallo, A., Ranaldi, F., Falco, V., Pasquali, V., Piro, G., Mita, G., and Di Sansebastiano, G. P. 2017. Vessel occlusion in three cultivars of Olea europaea naturally exposed to Xylella fastidiosa in open field. J. Phytopathol. 165:589–594

De Micco, V., Balzano, A., Wheeler, E. A., and Baas, P. 2016. Tyloses and Gums: A Review of Structure, Function and Occurrence of Vessel Occlusions. IAWA J. 37:186–205

Denny, T. P., Carney, B. F., Schell, M. A., and Others. 1990. Inactivation of multiple virulence genes reduces the ability of Pseudomonas solanacearum to cause wilt symptoms. Mol. Plant-Microbe Interact. 3

Earles, J. M., Knipfer, T., Tixier, A., Orozco, J., Reyes, C., Zwieniecki, M. A., Brodersen, C. R., and McElrone, A. J. 2018. In vivo quantification of plant starch reserves at micrometer resolution using X-ray microCT imaging and machine learning. New Phytol. 218:1260–1269

Fradin, E. F., and Thomma, B. P. 2006. Physiology and molecular aspects of Verticillium wilt diseases caused by V. dahliae and V. albo-atrum. Mol. Plant Pathol. 7:71–86

García, R. O., Kerns, J. P., and Thiessen, L. 2019. Ralstonia solanacearum Species Complex: A Quick Diagnostic Guide. Plant Health Prog. 20:7–13

Gluck-Thaler, E., Cerutti, A., Perez-Quintero, A., Butchacas, J., Roman-Reyna, V., Madhaven, V. N., Shantharaj, D., Merfa, M. V., Pesce, C., Jauneau, A., Vancheva, T., Lang, J. M., Allen, C., Verdier, V., Gagnevin, L., Szurek, B., Cunnac, S., Beckham, G., de la Fuente, L., Patel, H. K., Sonti, R. V., Bragard, C., Leach, J. E., Noël, L. D., Slot, J. C., Koebnik, R., and Jacobs, J. M. 2020. Repeated gain and loss of a single gene modulates the evolution of vascular pathogen lifestyles. Cold Spring Harbor Laboratory. :2020.04.24.058529

Grimault, V., Gélie, B., Lemattre, M., Prior, P., and Schmit, J. 1994. Comparative histology of resistant and susceptible tomato cultivars infected by Pseudomonas solanacearum. Physiol. Mol. Plant Pathol. 44:105–123

Ingel, B., Jeske, D. R., Sun, Q., Grosskopf, J., and Roper, M. C. 2019. Xylella fastidiosa endoglucanases mediate the rate of Pierce’s Disease development in Vitis vinifera in a cultivar-dependent manner. Mol. Plant. Microbe. Interact. 32:1402–1414

Ingel, B., Reyes, C., Massonnet, M., Boudreau, B., Sun, Y., Sun, Q., McElrone, A. J., Cantu, D., and Roper, M. C. 2020. Xylella fastidiosa causes transcriptional shifts that precede tylose formation and starch depletion in xylem. Mol. Plant Pathol.

Kaack, L., Altaner, C. M., Carmesin, C., Diaz, A., Holler, M., Kranz, C., Neusser, G., Odstrcil, M., Jochen Schenk, H., Schmidt, V., Weber, M., Zhang, Y., and Jansen, S. 2019. Function and three-dimensional structure of intervessel pit membranes in angiosperms: a review. IAWA J. 40:673–702

Kashyap, A., Planas-Marquès, M., Capellades, M., Valls, M., and Coll, N. S. 2020. Blocking intruders: inducible physico-chemical barriers against plant vascular wilt pathogens. J. Exp. Bot.

Khokhani, D., Tuan, T., Lowe-Power, T., and Allen, C. 2018. Plant assays for quantifying Ralstonia solanacearum virulence. Bio Protoc. 8

Kim, T.-H., Böhmer, M., Hu, H., Nishimura, N., and Schroeder, J. I. 2010. Guard cell signal transduction network: advances in understanding abscisic acid, CO2, and Ca2+ signaling. Annu. Rev. Plant Biol. 61:561–591

Knipfer, T., Brodersen, C. R., Zedan, A., Kluepfel, D. A., and McElrone, A. J. 2015. Patterns of drought-induced embolism formation and spread in living walnut saplings visualized using X-ray microtomography. Tree Physiol. 35:744–755

Li, M., Klein, L. L., Duncan, K. E., Jiang, N., Chitwood, D. H., Londo, J. P., Miller, A. J., and Topp, C. N. 2019. Characterizing 3D inflorescence architecture in grapevine using X-ray imaging and advanced morphometrics: implications for understanding cluster density. J. Exp. Bot. 70:6261–6276

Liu, H., Zhang, S., Schell, M. A., and Denny, T. P. 2005. Pyramiding unmarked deletions in Ralstonia solanacearum shows that secreted proteins in addition to plant cell-wall-degrading enzymes contribute to virulence. Mol. Plant. Microbe. Interact. 18:1296–1305

Lorenzini, G., Guidi, L., Nali, C., Ciompi, S., and Soldatini, G. F. 1997. Photosynthetic response of tomato plants to vascular wilt diseases. Plant Sci. 124:143–152

Lowe-Power, T., and Chipman, K. 2020. A meta-analysis of the known global distribution and host range of the Ralstonia species complex. Cold Spring Harbor Laboratory. :2020.07.13.189936

Lowe-Power, T. M., Hendrich, C. G., von Roepenack-Lahaye, E., Li, B., Wu, D., Mitra, R., Dalsing, B. L., Ricca, P., Naidoo, J., Cook, D., Jancewicz, A., Masson, P., Thomma, B., Lahaye, T., Michael, A. J., and Allen, C. 2018. Metabolomics of tomato xylem sap during bacterial wilt reveals Ralstonia solanacearum produces abundant putrescine, a metabolite that accelerates wilt disease. Environmental microbiology. 20:1330–1349

Lowe-Power, T. M., Khokhani, D., and Allen, C. 2018. How Ralstonia solanacearum exploits and thrives in the flowing plant xylem environment. Trends Microbiol. 26:929–942

McElrone, A. J., Choat, B., Parkinson, D. Y., MacDowell, A. A., and Brodersen, C. R. 2013. Using high resolution computed tomography to visualize the three dimensional structure and function of plant vasculature. J. Vis. Exp.

McElrone, A. J., Jackson, S., and Habdas, P. 2008. Hydraulic disruption and passive migration by a bacterial pathogen in oak tree xylem. J. Exp. Bot. 59:2649–2657

McGarvey, J. A., Denny, T. P., and Schell, M. A. 1999. Spatial-temporal and quantitative analysis of growth and EPS I production by Ralstonia solanacearum in resistant and susceptible tomato cultivars. Phytopathology. 89:1233–1239

Morella, N. M., Yang, S. C., Hernandez, C. A., and Koskella, B. 2018. Rapid quantification of bacteriophages and their bacterial hosts in vitro and in vivo using droplet digital PCR. J. Virol. Methods. 259:18–24

Nardini, A., Lo Gullo, M. A., and Salleo, S. 2011. Refilling embolized xylem conduits: is it a matter of phloem unloading? Plant Sci. 180:604–611

Opina, N., Tavner, F., Hollway, G., Wang, J. F., Li, T. H., Maghirang, R., Fegan, M., Hayward, A., Krishnapillai, V., Hong, W., and Others. 1997. A novel method for development of species and strain-specific DNA proves and PCR primers for identifying Burkholderia solanacearum (formerly Pseudomonas solanacearum). Asia-pacific Journal of Molecular Biology and Biotechnology. 5:19–30

Pérez-Donoso, A. G., Lenhof, J. J., Pinney, K., and Labavitch, J. M. 2016. Vessel embolism and tyloses in early stages of Pierce’s disease. Aust. J. Grape Wine Res. 22:81–86

Pérez-Donoso, A. G., Sun, Q., Roper, M. C., Greve, L. C., Kirkpatrick, B., and Labavitch, J. M. 2010. Cell wall-degrading enzymes enlarge the pore size of intervessel pit membranes in healthy and Xylella fastidiosa-infected grapevines. Plant Physiol. 152:1748–1759

Planas-Marquès, M., Kressin, J. P., Kashyap, A., Panthee, D. R., Louws, F. J., Coll, N. S., and Valls, M. 2020. Four bottlenecks restrict colonization and invasion by the pathogen Ralstonia solanacearum in resistant tomato. J. Exp. Bot. 71:2157–2171

Prior, P., Ailloud, F., Dalsing, B. L., Remenant, B., Sanchez, B., and Allen, C. 2016. Genomic and proteomic evidence supporting the division of the plant pathogen Ralstonia solanacearum into three species. BMC Genomics. 17:90

Rahman, M. A., Abdullah, H., and Vanhaecke, M. 1999. Histopathology of susceptible and resistant Capsicum annuum cultivars infected with Ralstonia solanacearum. J. Phytopathol. 147:129–140

Rajagopal, V., Sumathykuttyamma, B., and Patil, K. D. 1987. Water relations of coconut palms affected with root (wilt) disease. New Phytol. 105:289–293

Roper, M. C., Greve, L. C., Warren, J. G., Labavitch, J. M., and Kirkpatrick, B. C. 2007. Xylella fastidiosa requires polygalacturonase for colonization and pathogenicity in Vitis vinifera grapevines. Mol. Plant. Microbe. Interact. 20:411–419

Schwarze, F. W., and Landmesser, H. 2000. Preferential degradation of pit membranes within tracheids by the basidiomycete Physisporinus vitreus. Holzforschung. 54:461–462

Sperry, J. S., and Hacke, U. G. 2004. Analysis of circular bordered pit function I. Angiosperm vessels with homogenous pit membranes. Am. J. Bot. 91:369–385

Stevenson, J. F., Matthews, M. A., Carl Greve, L., Labavitch, J. M., and Rost, T. L. 2004. Grapevine susceptibility to Pierce’s Disease II: progression of anatomical symptoms. Am. J. Enol. Vitic. 55:238–245

Sun, Q., Greve, L. C., and Labavitch, J. M. 2011. Polysaccharide compositions of intervessel pit membranes contribute to Pierce’s disease resistance of grapevines. Plant Physiol. 155:1976–1987

Sun, Q., Sun, Y., Walker, M. A., and Labavitch, J. M. 2013. Vascular occlusions in grapevines with Pierce’s disease make disease symptom development worse. Plant Physiol. 161:1529–1541

Tyree, M. T., and Sperry, J. S. 1989. Vulnerability of xylem to cavitation and embolism. Annu. Rev. Plant Physiol. Plant Mol. Biol. 40:19–36

Tyree, M. T., and Zimmermann, M. H. 2002. Hydraulic architecture of whole plants and plant performance. Pages 175–214 in: Xylem Structure and the Ascent of Sap, M.T. Tyree and M.H. Zimmermann, eds. Springer Berlin Heidelberg, Berlin, Heidelberg.

Vasse, J., Frey, P., and Trigalet, A. 1995. Microscopic studies of intercellular infection and protoxylem invasion of tomato roots by Pseudomonas solanacearum. Mol. Plant. Microbe. Interact. 8:241–251

Weiss, R. F. 1970. The solubility of nitrogen, oxygen and argon in water and seawater. Deep Sea Research and Oceanographic Abstracts. 17:721–735

Windt, C. W., Vergeldt, F. J., De Jager, P. A., and Van As, H. 2006. MRI of long-distance water transport: a comparison of the phloem and xylem flow characteristics and dynamics in poplar, castor bean, tomato and tobacco. Plant Cell Environ. 29:1715–1729

Yadeta, K. A., and Thomma, B. P. H. J. 2013. The xylem as battleground for plant hosts and vascular wilt pathogens. Front. Plant Sci. 4:97

